# Cellular co-infection can modulate the efficiency of influenza A virus production and shape the interferon response

**DOI:** 10.1101/752329

**Authors:** Brigitte E. Martin, Jeremy D. Harris, Jiayi Sun, Katia Koelle, Christopher B. Brooke

## Abstract

During viral infection, the numbers of virions infecting individual cells can vary significantly over time and space. The functional consequences of this variation in cellular multiplicity of infection (MOI) remain poorly understood. Here, we rigorously quantify the phenotypic consequences of cellular MOI during influenza A virus (IAV) infection over a single round of replication in terms of cell death rates, viral output kinetics, interferon and antiviral effector gene transcription, and superinfection potential. By statistically fitting mathematical models to our data, we precisely define specific functional forms that quantitatively describe the modulation of these phenotypes by MOI at the single cell level. To determine the generality of these functional forms, we compare two distinct cell lines (MDCK cells and A549 cells), both infected with the H1N1 strain A/Puerto Rico/8/1934 (PR8). We find that a model assuming that infected cell death rates are independent of cellular MOI best fits the experimental data in both cell lines. We further observe that a model in which the rate and efficiency of virus production increase with cellular co-infection best fits our observations in MDCK cells, but not in A549 cells. In A549 cells, we also find that induction of type III interferon, but not type I interferon, is highly dependent on cellular MOI, especially at early timepoints. This finding identifies a role for cellular co-infection in shaping the innate immune response to IAV infection. Finally, we show that higher cellular MOI is associated with more potent superinfection exclusion, thus limiting the total number of virions capable of infecting a cell. Overall, this study suggests that the extent of cellular co-infection by influenza viruses may be a critical determinant of both viral production kinetics and cellular infection outcomes in a host cell type-dependent manner.

**AUTHOR SUMMARY:** During influenza A virus (IAV) infection, the number of virions to enter individual cells can be highly variable. Cellular co-infection appears to be common and plays an essential role in facilitating reassortment for IAV, yet little is known about how cellular co-infection influences infection outcomes at the cellular level. Here, we combine quantitative *in vitro* infection experiments with statistical model fitting to precisely define the phenotypic consequences of cellular co-infection in two cell lines. We reveal that cellular co-infection can increase and accelerate the efficiency of IAV production in a cell line-dependent fashion, identifying it as a potential determinant of viral replication kinetics. We also show that induction of type III, but not type I, interferon is highly dependent upon the number of virions that infect a given cell, implicating cellular co-infection as an important determinant of the host innate immune response to infection. Altogether, our findings show that cellular co-infection plays a crucial role in determining infection outcome. The integration of experimental and statistical modeling approaches detailed here represents a significant advance in the quantitative study of influenza virus infection and should aid ongoing efforts focused on the construction of mathematical models of IAV infection.

## INTRODUCTION

Cellular co-infection plays an important, yet poorly defined, role in shaping the outcome of influenza A virus (IAV) infection. By facilitating reassortment between incoming viral genomes, cellular co-infection can give rise to new viral genotypes with increased fitness or emergence potential (1). Cellular co-infection can also enhance the replicative potential of the virus by promoting the complementation and multiplicity reactivation of the semi-infectious particles that constitute the bulk of influenza virus populations (2–4). Despite its clear importance, the prevalence and the specific functional consequences of cellular co-infection during IAV infection remain largely unknown.

We and others have previously shown that cellular co-infection can be common *in vivo* (5–7). There is growing evidence that IAV replication and spread is focal and thus that the distribution of individual virions across cells and tissues is highly spatially structured, resulting in foci of high cellular multiplicity of infection (MOI) (8–11). Given the dynamic distribution of virions over time and space during infection, it is likely that the MOIs of individual infected cells are highly variable. This raises the question of whether variation in the number of virions that infect a cell has distinct phenotypic consequences. If so, it could have significant implications for understanding IAV infection dynamics as two viral populations of identical size and genome sequence could give rise to divergent infection outcomes if the dispersal patterns of virions (and thus the MOI distribution across cells) differs.

Several previous studies have suggested that the number of virions that enter a given cell (referred to throughout as “viral input” or “cellular MOI”) may affect replication kinetics and interferon (IFN) induction (12–16). However, the phenotypic consequences of cellular MOI during IAV infection have not yet been rigorously or comprehensively quantified. In this manuscript, we focus on the infection dynamics of two cell lines (MDCK and A549) infected with A/Puerto Rico/8/1934 (PR8). We combine precise single-cycle infection experiments with statistical model fitting to reveal that cellular MOI can significantly alter virus production rates, the host transcriptional response to infection, and the potential for superinfection. In doing so, we precisely define functional forms that can account for the observed relationships between viral input and the phenotypes of infected cells, information that will aid future efforts to quantitatively model IAV infection. Altogether, these results reveal and define an underappreciated role for cellular co-infection in shaping the outcome of IAV infection.

## RESULTS

To define how variation in cellular MOI affects viral replication dynamics and the host response to infection, we infected either MDCK or A549 cells with PR8 across a 100-fold range of bulk MOIs. The working stock of PR8 that we used has a physical particle (matrix segment genome equivalents (GE)/mL) to fully infectious particle ratio (tissue culture infectious dose 50 (TCID50)/mL) of 8.19, similar to previous reports (2). To eliminate the confounding effects of secondary spread within culture on infectious outcomes, we limited infections to a single cycle by treating cells with 25 mM NH4Cl at 2 hpi (17). Thus, in all experiments described below, we were examining outcomes following a single round of infection.

### Precise quantification of the actual bulk MOI

An accurate assessment of the phenotypic consequences for cellular co-infection depends upon the precise measurement of the average number of viral genomes that actually contribute to viral replication and/or immune activation. We suspected that the standard method for calculating MOI in bulk cell culture, based on the dilutions of viral working stock used, may overestimate the actual number of virions that successfully infect due to incomplete virion adsorption or entry.

To calculate the “actual” bulk MOI that contributed to infection, we quantified the fraction of virus inoculum that was successfully bound or taken up during the adsorption phase for both MDCK and A549 cells. We first measured the concentration of virus present in the inoculum added to cells (0 hpi) and the amount of remaining unbound virus following 1-hour adsorption at 4°C (1 hpi) across a range of intended bulk MOIs (**Fig 1A**) in triplicate by RT-qPCR. Cells were washed extensively and transitioned to growth media following the 1 hr adsorption period. To confirm that our post-adsorption washes effectively removed any unbound inoculum virus that could artificially inflate subsequent viral output measurements, we quantified extracellular virus immediately following wash and found that viral loads were negligible (<10^3^ genome equivalent (GE)/mL; data not shown). At 2 hours post-adsorption (3 hpi), we again measured the amount of extracellular virus present and found titers that were surprisingly high for this early timepoint, especially at an intended bulk MOI of 10 (**Fig 1A**).

**Figure 1.**
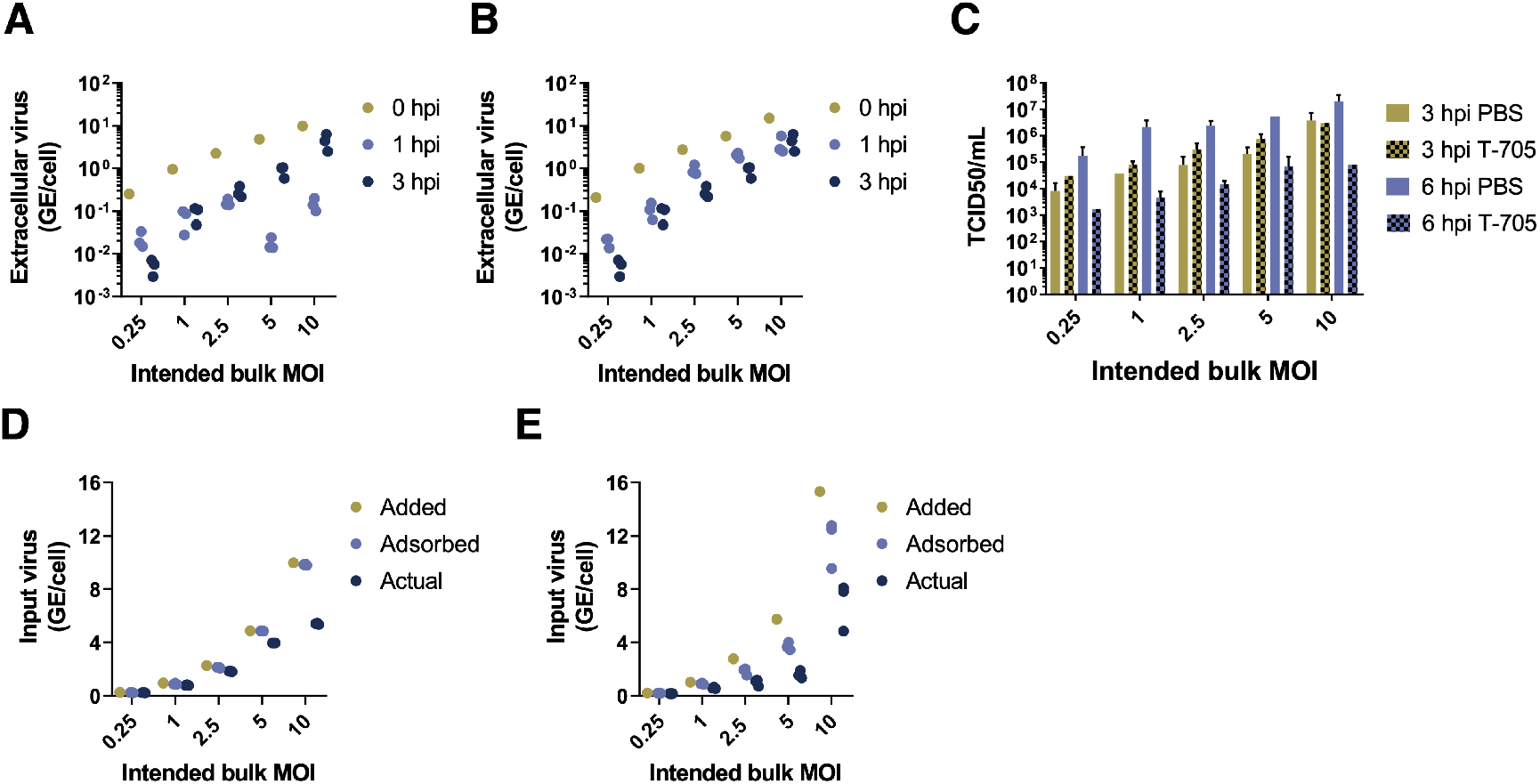
Precise quantification of actual bulk MOI. **(A)** Extracellular virus concentrations present in the inoculum (0 hpi), remaining unbound following adsorption (1 hpi), and measured extracellularly at 3 hpi for both MDCK and A549 cells, at the indicated bulk MOIs. Extracellular virus concentrations are given in units of viral genome equivalents (GE) per cell. 0 hpi data points show single viral measurements from the inoculum used for all replicates. 1 hpi and 3 hpi points show individual measurements from replicate infection wells. **(B)** Virus output at 3 or 6 hpi from MDCK cells infected in triplicate with PR8 at the indicated intended MOIs and treated with either PBS or with 40 uM T-705 for 2 hrs prior to infection and throughout the duration of infection. **(C)** Quantification by RT-qPCR of virus present in inoculum (0 hpi; “added MOI”), virus adsorbed into cells after 1 hpi (calculated by subtracting extracellular virus present at 1 hpi from that present in inoculum at 0 hpi; “adsorbed MOI”), and virus expected to actually contribute to infection (calculated by subtracting the average of three replicates of extracellular virus measurements at 3 hpi from adsorbed MOI; “actual MOI”) for MDCK and A549 cells.

To determine whether extracellular virus present at 3 hpi was newly produced or carried-over from the inoculum, we repeated the above experiments in MDCK cells treated from 2 hours prior to infection through the duration of infection (6 hpi) with 40 *μ*M of the antiviral drug T-705, which inhibits the production of viral progeny (18). Drug treatment significantly reduced viral titers at 6 hpi, but not 3 hpi, indicating that extracellular virus measured at 3 hpi consists of inoculum virus that was taken up but then released without actually infecting the cell (**Fig 1B**). Thus, to precisely quantify the actual bulk MOI of virus that contributed to infection in each sample, we subtracted the amounts of extracellular virus detected at both 1 hpi and 3 hpi (for this, we used the average of three experimental replicates since they were from independent samples) from our measurements of bulk MOI at 0 hpi. This gave us an actual bulk MOI range of 0.23-5.45 for MDCK cells and 0.16-8.09 for A549 cells (**Fig 1C**). Hereafter, we use these estimated actual bulk MOIs instead of intended bulk MOIs to quantify the effects of cellular MOI on virus production kinetics and cellular infection outcomes and refer to these actual bulk MOIs simply as ‘bulk MOIs’.

### Assumption of viral infection distribution

To quantify the effects of cellular MOI on virus production kinetics and cellular infection outcomes, an assumption needs to be made about how virus particles are distributed across cells. It is often assumed that the number of virions that infect a given cell follows a Poisson distribution with both the mean and variance in cellular MOI equal to the overall bulk MOI. However, empirical support for this assumption is lacking. To accommodate potential deviations from the Poisson distribution, we instead assume that virions are distributed across cells according to a negative binomial distribution. This distribution allows for virions to be overdispersed when the dispersion parameter is small, with the variance in cellular MOI across cells exceeding the mean cellular MOI. That is, in the case of overdispersion (relative to a Poisson assumption), a higher fraction of cells will have either low/no viral input or high viral input relative. A negative binomial distribution, however, also allows for virions to be Poisson-distributed across cells when the dispersion parameter is large. As such, this distribution provides us with a more flexible approach for modeling the distribution of viral particles across cells. To be thorough, we also considered the possibility of virions being distributed according to a zero-inflated Poisson distribution, where a fraction of cells remain uninfected, and the remaining cells have cellular multiplicities of infection governed by a Poisson distribution with a higher mean to account for the cells that remain uninfected. Similar to the negative binomial distribution, the zero-inflated Poisson distribution allows for the incorporation of cellular heterogeneity in a phenomenological manner.

When fitting the cell death models (described in the next section) to experimental data, we simultaneously estimate parameters of these viral infection distribution models using fluorescence-activated cell sorting (FACS) measurements taken at 18 hpi (**Fig S1**). When fitting the remaining statistical models (described below), we take as given the most well-supported, already-parameterized viral infection distribution and the most well-supported, already-parameterized cell death model.

### Cell death rates are time-dependent but virus input-independent in both MDCK and A549 cells

The overall productivity of an infected cell depends in part upon how long the cell survives following infection, and we hypothesized that cellular lifespans might be affected by cellular MOI. We quantified how differences in bulk MOI affected cell survival at different times post-infection (compared with mock infected cells) using trypan blue exclusion. As expected, the number of surviving MDCK cells generally decreased with higher bulk MOIs for each timepoint tested (**Fig 2**). To determine whether this observed decrease in cell survival was simply due to increases in the fraction of infected cells, or if, instead, cells co-infected with multiple virions died at faster rates than singly infected cells, we statistically fit a set of mathematical models to the experimental data (Methods).

**Figure 2.**
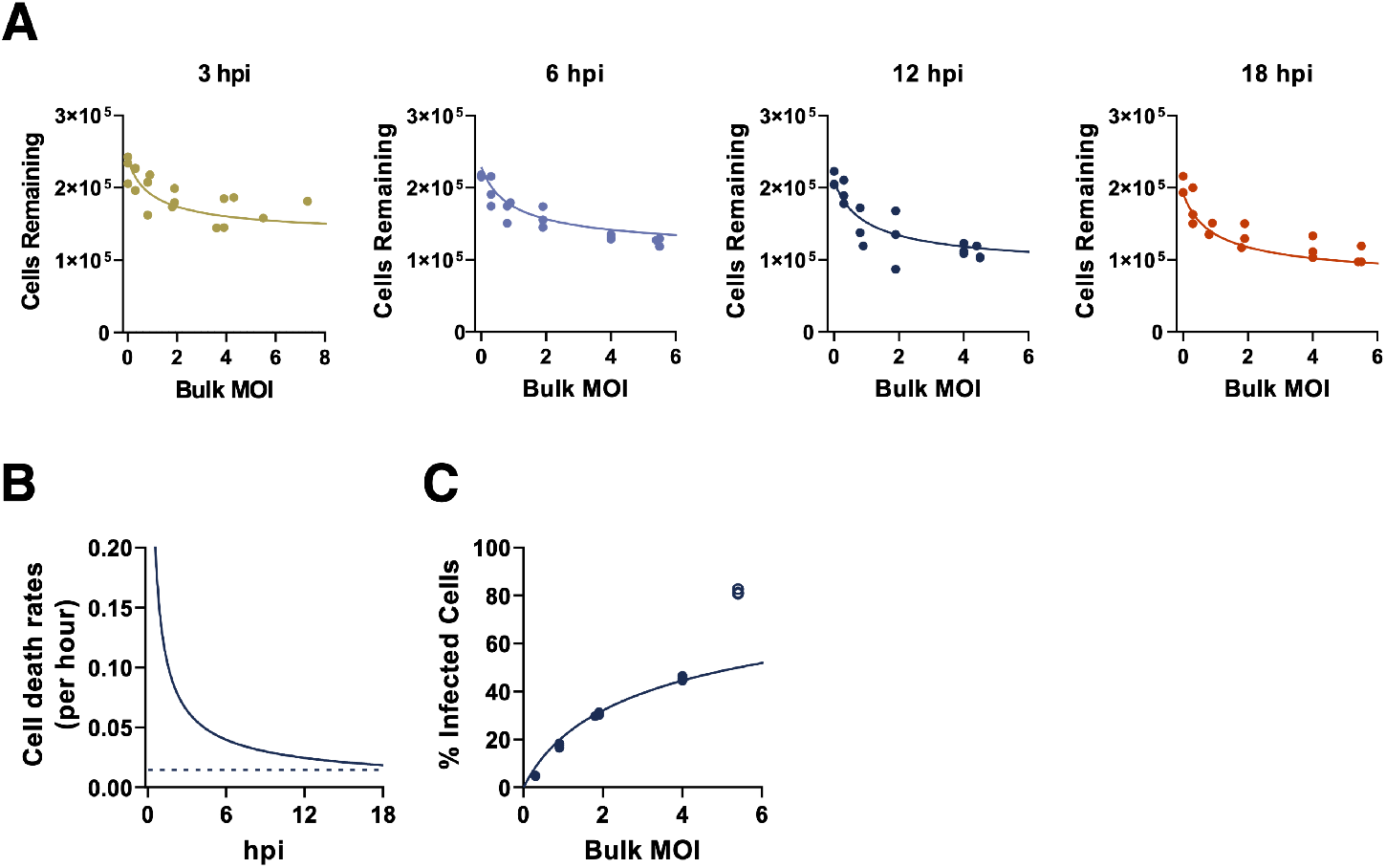
MDCK cell death rates are time-dependent and input-independent. **(A)** The numbers of MDCK cells surviving following infection, as determined by by trypan blue exclusion at the indicated timepoints across our experimental range of bulk MOIs. Values represent the number of trypan blue negative cells in each sample at 3, 6, 12, and 18 hours post infection (hpi). Lines indicate the best model fit to these data, which is given by the time-dependent, input-independent cell death rate model, as parameterized in **Table S1. (B)** Estimated input-independent cell death rate over the course of cellular infection (solid) and constant background cell death rate (dashed; obtained from mock infected cells). **(C)** Percent of surviving MDCK cells that are infected at 18 hpi, as measured by FACS (FACS plots shown in **Fig S1**). The line indicates the negative binomial distribution with time-dependent input-independent cell death rate model fit evaluated at 18 hpi. Statistical parameterization of this model (overdispersion parameter *r*= 0.597; **Table S1**) indicates a high level of overdispersion and significant deviation from a Poisson-distributed model. Data from the highest bulk MOI were excluded from model fits due to the lack of confidence in the accuracy of FACS measurements at the highest MOI examined.

We first considered a model in which all infected cells died at a constant (and identical) rate over the course of cellular infection, under all three possible viral infection distributions (Poisson, negative binomial, and zero-inflated Poisson). Jointly fitting this cell death model and the viral infection distribution to the FACS data and the number of surviving MDCK cells, we found that the negative binomial distribution was most well-supported by the data (**Table S1**; **Fig S2B**), indicating that the distribution of virions across cells is highly overdispersed. This model, with a constant and identical (input-independent) cell death rate, however systematically overestimated the number of surviving cells at early time points following infection (3 and 6 hpi; **Fig S2A**) and systematically underestimated the number of surviving cells at the latest time point (18 hpi; **Fig S2A**). We next considered a model in which infected cell death rates were allowed to vary over the course of infection (i.e., death rates could be time-dependent), with rates that were still independent of cellular MOI (i.e., death rates were input-independent). For this model, we chose a Weibull hazard function as the functional form to describe the time-varying death rate because of its common use in survival analysis. Jointly fitting this model and the viral infection distributions to the MDCK cell data and the FACS data yielded, for all three viral infection distributions considered, an infected cell death rate that decreased over the course of cellular infection (**Table S1**). Of the viral infection distribution models, the negative binomial one was again most well-supported by the data (**Table S1**). Under this distribution, the parameterized time-dependent, input-independent cell death model did not over- or underestimate the number of surviving cells at any of the four time points considered (**Fig 2A**) and was able to quantitatively reproduce the measured FACS data (**Fig 2C**). This time-dependent cell death rate model was statistically preferred over the time-independent cell death rate model (**Table S1**).

We next considered a time-independent, input-dependent model of cell death rates in which the death rate of a given cell increased with viral input. Again, the negative binomial distribution was preferred over both the Poisson model and the zero-inflated Poisson model (**Table S1**). Under this negative binomial model for the viral infectivity distribution, the cell death model yielded similar fits as the time-independent, input-independent model (**Fig S3**). Indeed, its statistical parameterization effectively reduced this model to the time-independent, input-independent model (**Table S1**). This model had less statistical support than the time-independent, input-independent model though because of its higher complexity (**Table S1**). Finally, we considered a general time-dependent, input-dependent model that could reduce, under specific parameterizations, to either a time-independent model, an input-independent model, or both (**Table S1**). Again, the negative binomial distribution was preferred over both the Poisson model and the zero-inflated Poisson model (**Table S1**). Under this negative binomial model, statistical parameterization of this cell death model yielded fits similar to those of the time-dependent, input-independent model. Indeed, the parameterization of this model effectively reduced this model to this simpler, preferred model. Because of its greater complexity, this cell death rate model had less statistical support than the time-dependent, input-independent model. Overall, we found that the time-dependent, input-independent cell death rate model was the most well-supported cell death rate model for the MDCK cell data. This finding indicates that lower levels of cell survival at higher bulk MOIs (as observed in Figure 2) are likely simply due to there being a higher fraction of cells that are infected at higher MOIs, rather than being due to cell death rates being higher in cells with higher viral input.

For all four models considered, the fits to the FACS data indicated that viral particles are decidedly not Poisson-distributed across cells in the bulk MOI range considered. Instead, the statistical parameterizations of all four models point towards a high level of viral overdispersion (**Table S1**). Models that forced the assumption of a Poisson distribution were unable to quantitatively reproduce the FACS data (**Fig S4A**). Models that allowed for a more flexible negative binomial distribution were statistically strongly preferred over those that assumed a Poisson distribution of virions across cells (**Fig S4A**, **Table S1**). The negative binomial model was also preferred over the zero-inflated Poisson model (**Table S1**), although not as strongly as over the standard Poisson model.

For A549 cells, we considered the same set of cell death rate models and viral infectivity distribution models as we did for MDCK cells. Here, we again found that the best model fits assumed a negative binomial distribution for viral infectivity and that the time-dependent, input-independent cell death rate model was statistically preferred over the other three models (**Table S2**; **Fig 3**). In sum, by fitting to FACS data and a time course of surviving cell numbers, we found that a model that assumes that infected cell death rates are changing with time since infection (but not with viral input) is most well-supported by both MDCK and A549 data. In both cell lines, the parameterized model results in a cellular death rate that declines with time since infection. This finding, although not intuitive, is visually supported by the data. For example, it is visually apparent that in the MDCK cell line, at a given bulk MOI, that the number of surviving cells rapidly declines until 6 hpi, and then stays at similar levels until 18 hpi.

**Figure 3.**
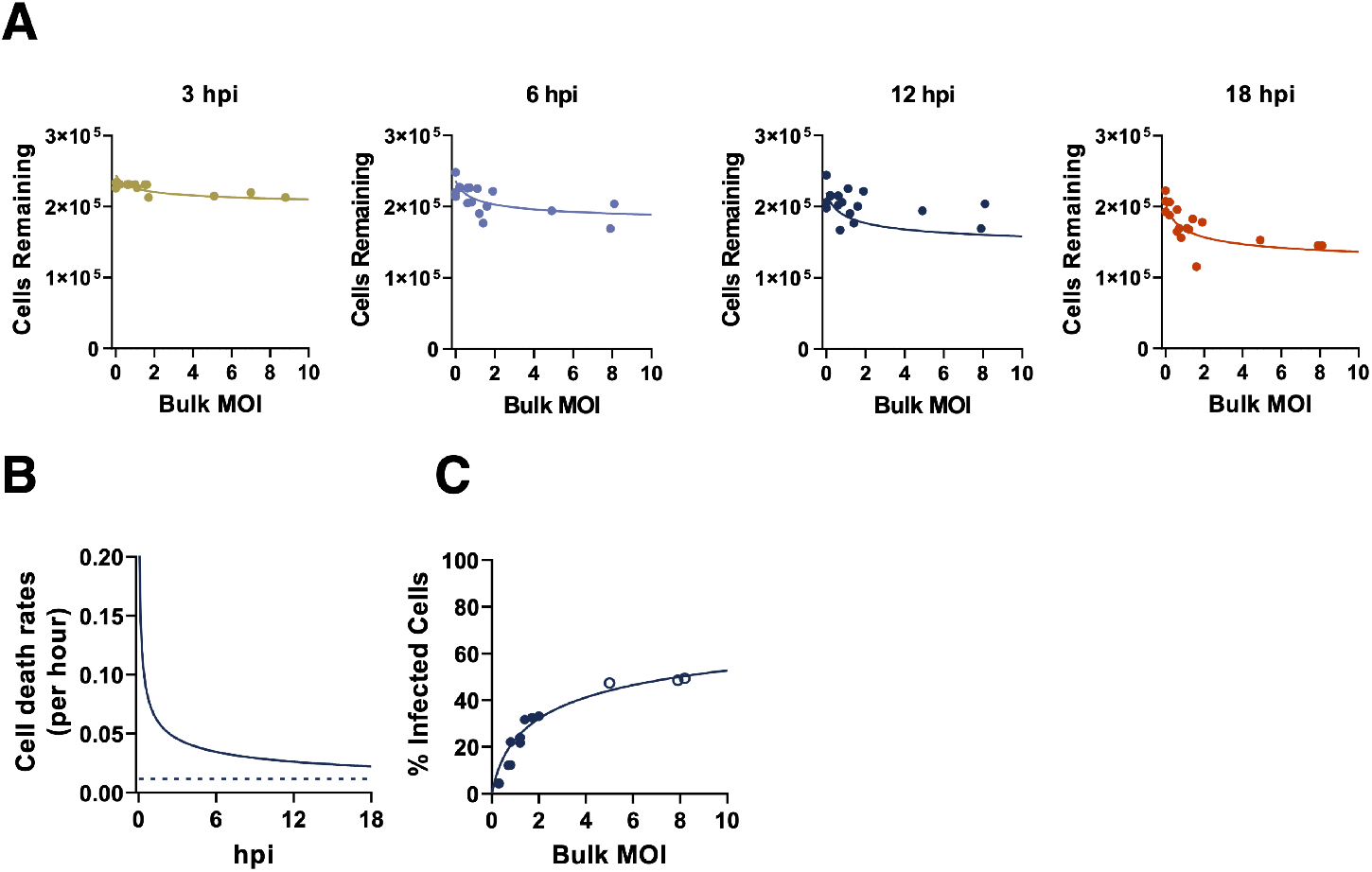
A549 cell death rates are time-dependent and input-independent. **(A)** The numbers of A549 cells surviving following infection, as determined by by trypan blue exclusion at the indicated timepoints across our experimental range of bulk MOIs. Values represent the number of trypan blue negative cells in each sample. Lines indicate the best model fit to these data, given by the time-dependent, input-independent cell death rate model, as parameterized in **Table S2**. **(B)** Estimated A549 cell death rate (solid) and the constant background cell death model (dashed), fitting to both mock infected cells and MOI treatments over the course of infection. **(C)** Percent of surviving A549 cells that are infected at 18 hpi (FACS data). The line indicates the negative binomial distribution (*r*= 0.338; **Table S2)** with time-dependent input-independent cell death rate model fit evaluated at 18 hpi. Data from the highest bulk MOI were excluded from model fits due to the lack of confidence in the accuracy of FACS measurements at the highest MOI examined.

### Cellular co-infection increases the rate of virus production in MDCK cells but not in A549 cells

We next asked how changes in bulk MOI affected virus production over a single cycle of infection in both MDCK and A549 cells. In the same experiments described above, we measured the total viral output (in GE/mL) from cells infected at different bulk MOIs at 6, 12, and 18 hpi. Not surprisingly, we observed that viral output from both cell lines was significantly increased at higher bulk MOIs for all time points tested (**Fig 4A**).

**Figure 4.**
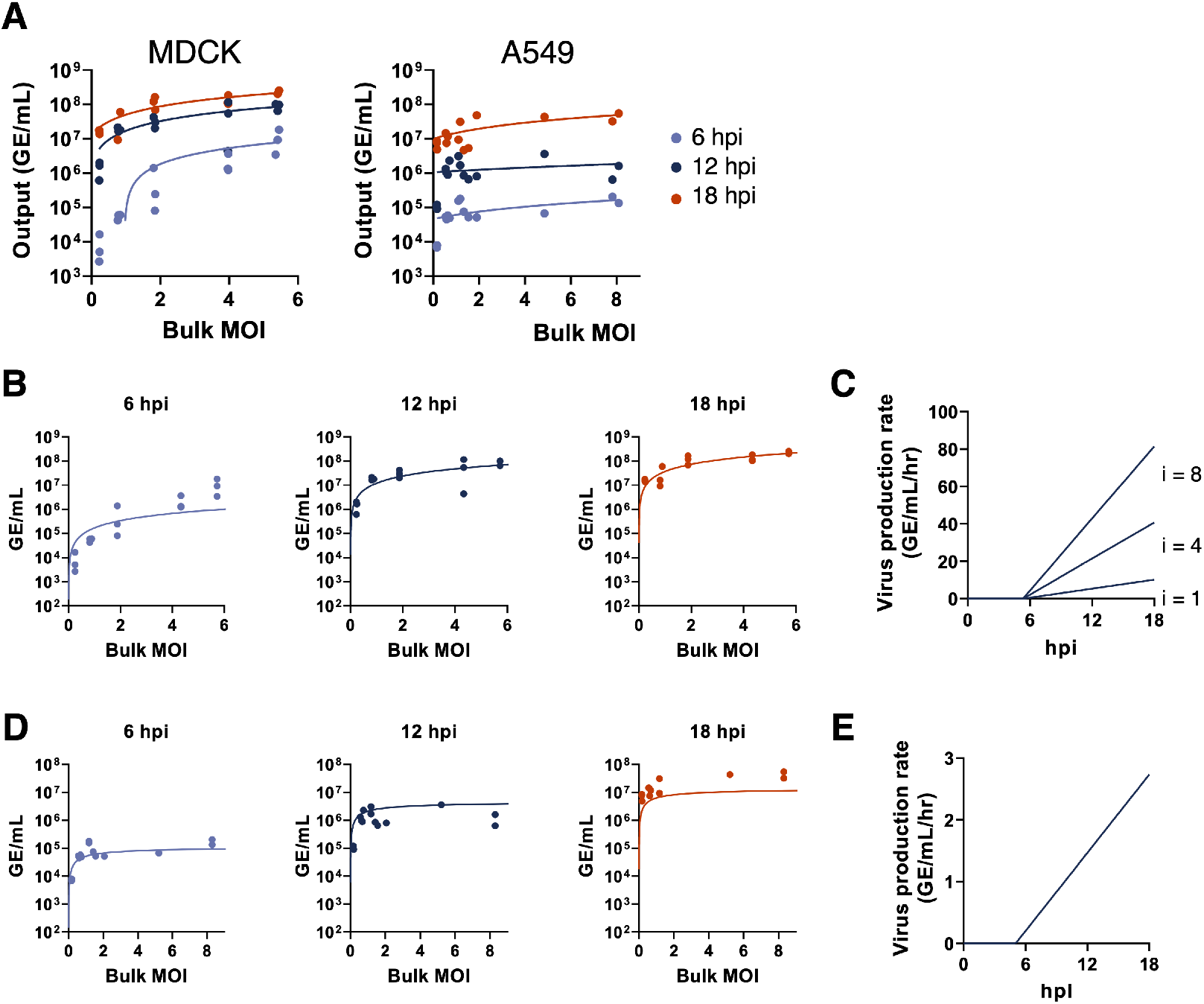
Cellular co-infection increases the rate of virus production in MDCK cells but not in A549 cells. **(A)** Viral output from single cycle infections of MDCK cells or A549 cells with PR8 over a range of bulk MOIs, as measured by RT-qPCR at the indicated hours post infection. Each data point represents both the actual MOI and viral output from a single infection well. Trend lines show linear regressions performed on the log-log scale. All regression slopes for MDCK cells are significantly positive (p < 0.0001; 6 hpi: slope = 2.2, R^2^ = 0.90; 12 hpi: slope = 1.15, R^2^ = 0.71; 18 hpi: slope = 0.88, R^2^ = 0.78). All regression slopes for A549 cells are significantly positive (p < 0.01; 6 hpi: slope = 0.71, R^2^ = 0.68; 12 hpi: slope = 0.61, R^2^ = 0.44; 18 hpi: slope = 0.48, R^2^ = 0.49). (B) The time-dependent, linear input-dependent model of virus production fit to virus output data from MDCK cells at the indicated timepoints. (C) Visualization of the time-dependent, linear input-dependent model of virus production from infected MDCK cells, parameterized with values shown in Table S3. Rates of virus production are shown over time for cells with virus input of ? = 1,4, and 8. (D). Same as panel (B) but for A549 cells. (E) Visualization of the time-dependent, input-independent model of virus production rate from infected A549 cells.

These findings raised the question of whether the increase in viral output at higher bulk MOIs is simply due to higher numbers of infected cells at higher bulk MOIs, or whether individual infected cells produce more virus output when co-infected by multiple virions. To infer the functional relationship between cellular MOI and viral output, we statistically fit several mathematical models of virus production to our data (Methods). Each of these models incorporated our results of cell death kinetics, described in the previous section. They each also incorporated the negative binomial distribution for viral infectivity, described in the previous section. The incorporation of these models was needed to accommodate the fact that only surviving infected cells should have the ability to contribute to viral output. For both MDCK and A549 cell lines, the first three models we fit were time-independent with respect to the rate of virus production but differed in their dependence upon cellular input. The statistical fits of these time-independent models to the data are summarized in **Table S3** (MDCK cells) and **Table S4** (A549 cells). In both cell lines, we found that even the best models overestimated virus output at 6 hpi and underestimated virus output at 18 hpi, across all bulk MOIs examined (**Fig S5**).

These results suggest that the rate of virus production increases over time. Therefore, we next considered a set of time-dependent models of virus production rate. Specifically, in these models we allowed for a time delay in virus production to account for the eclipse phase of infection. Following this delay, the rate of virus production was assumed to depend linearly on time. Input-dependency on the rate of virus production was incorporated differently by the three different time-dependent models. The first time-dependent model assumed that the rate of virus production was input-independent but increased linearly in time following the initial delay in virus production. When fit to the MDCK cell data, this model underestimated the amount of virus produced at high MOI values (**Fig S6**). We next considered a linear input-dependent model in which the rate of virus production (at any given point in time following the delay in virus production) increases linearly with the number of viruses infecting a given cell. In MDCK cells, this time-dependent model with linear input-dependent virus production rate was more supported by the data than the input-independent model (**Table S3**). Lastly, we considered a saturating input-dependent model of virus production. In this model, at any given time point following the delay in virus production the rate of virus production increases with the cellular MOI, saturating at high levels of cellular MOI. The parameter estimates from fitting the model to MDCK virus production data gave similar model fits as the linear input-dependent model. However, due to the increased complexity of this model, the saturating input-dependent model had less statistical support than the linear input-dependent model (**Table S3**).

Of the models considered for MDCK cells, we find that the time-dependent model with linear input-dependent virus production rate is best supported by the data. This model captures virus production vs. bulk MOI well at 12 and 18 hpi, but we do note discrepancies at 6 hpi between the fit of this model and the experimental data (**Fig 4B**). At 6 hpi, virus output at low bulk MOI is lower than that predicted by the model, while virus output at high bulk MOI is higher than that predicted by the model. This indicates that the delay in virus production may be longer for cells with low virus input than with higher virus input. A model that allows for the onset times of viral production to depend on cellular MOI may therefore better capture the relationship between viral output and cellular MOI at this early time point. Our overall results are consistent with previous reports of earlier and higher rates of replication under high experimental MOI conditions (19,20).

For A549 cells, we considered the same set of time-dependent virus production rate models. In contrast to MDCK cells, we did not find that virus production rates depended on virus input; the input-independent model was most supported by the data (**Table S4**; **Fig 4D,E**). The linear input-dependent model both overestimated viral output at high bulk MOI values at 6 and 12 hpi and underestimated output at low bulk MOI values at 18 hpi (**Fig S7**). The saturating input-dependent model yielded a slightly better fit to the data than the input-independent model, but due to its increased model complexity had less statistical support (**Table S4**). The statistically preferred input-independent virus production model reproduces the experimental data well at 6 hpi; however, we note that the model slightly overestimates viral output at 12 hpi and underestimates the output at 18 hpi (**Fig 4D**). This suggests that the rate of virus production may level off over time, instead of continuing to increase as assumed by our model structure. Similar to MDCK cells, all models fit to A549 cell data indicate that the onset of virus production occurs at approximately 5 hpi (**Table S4**; **Fig 4E**). In contrast to MDCK cells, though, we find that the rates of virus production in A549 cells do not increase with cellular MOI, revealing that the relationship between cellular MOI and viral output is cell type-specific.

### Increased cellular co-infection enhances the efficiency of viral progeny production in MDCK cells but not in A549 cells

To control for the differences in input genomes across the different bulk MOI conditions, we calculated per capita virus production by dividing the number of output viral genomes by the number of input genomes (**Fig 5**). For MDCK cells, increasing the bulk MOI significantly increased per capita virus output at 6 hpi (R^2^ = 0.71, p < 0.0001). However, this positive correlation between bulk MOI and per capita virus output was no longer apparent at later timepoints (12 hpi p = 0.47; 18 hpi p = 0.38), likely due to the eventual saturation of virus production rates at high cellular MOIs. Importantly, MDCK cells infected at higher bulk MOI crossed the threshold into positive virus production (virus output/input > 1) by 6 hpi, while the lower MOI infections lagged. In contrast, increasing bulk MOI had no significant effects on per capita virus output from A549 cells at 6 hpi and 12 hpi (p = 0.11 and p = 0.06, respectively), and actually decreased per capita output at 18 hpi (R^2^ 0.54, p = 0.0029). These results suggest that increasing the number of virions introduced into a given cell can increase or potentially decrease the efficiency of viral progeny production in a cell-type specific manner.

**Figure 5.**
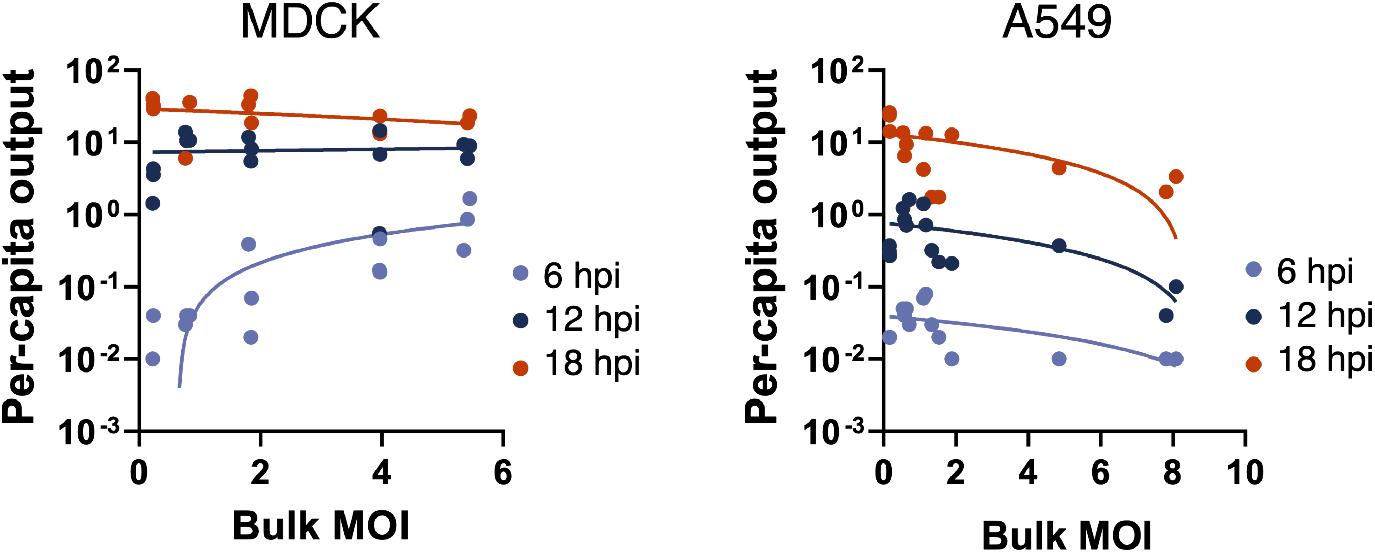
Cellular co-infection increases the efficiency of virus production in MDCK cells but not in A549 cells. Per capita virus output calculated as the ratio of output viral genomes over input viral genomes for individual infection replicates at the indicated timepoints vs. bulk MOI for MDCK or A549 cells. Trend lines represent linear regressions performed on the log-log scale for each time point plotted on the semilog scale. P values for the correlations between bulk MOI and per capita virus output at the indicated timepoints are shown.

### Cellular co-infection enhances ISG induction in A549 cells but not MDCK cells

We next asked whether cell type-specific differences in innate immune activation might explain the divergent effects of cellular MOI on virus output in MDCK and A549 cells. We examined the effects of increasing cellular MOI on the expression of two interferon stimulated genes (ISGs) known to inhibit IAV replication (ISG15 and Mx1) (**Fig 6**). In MDCK cells, we found no significant positive correlations between ISG expression levels and cellular MOI at either timepoint tested (**Fig 6A**). In A549 cells, expression of ISG15 was positively correlated with bulk MOI at 18 hpi, but not 8 hpi, while Mx1 expression levels were positively correlated with bulk MOI at both timepoints (**Fig 6B**). These results suggest that the benefits of increasing cellular MOI for viral progeny production that we observed in MDCK cells may be offset in A549 cells by increasing anti-viral ISG activity, thus potentially explaining the divergent effects of cellular MOI on virus output in these two cell types.

**Figure 6.**
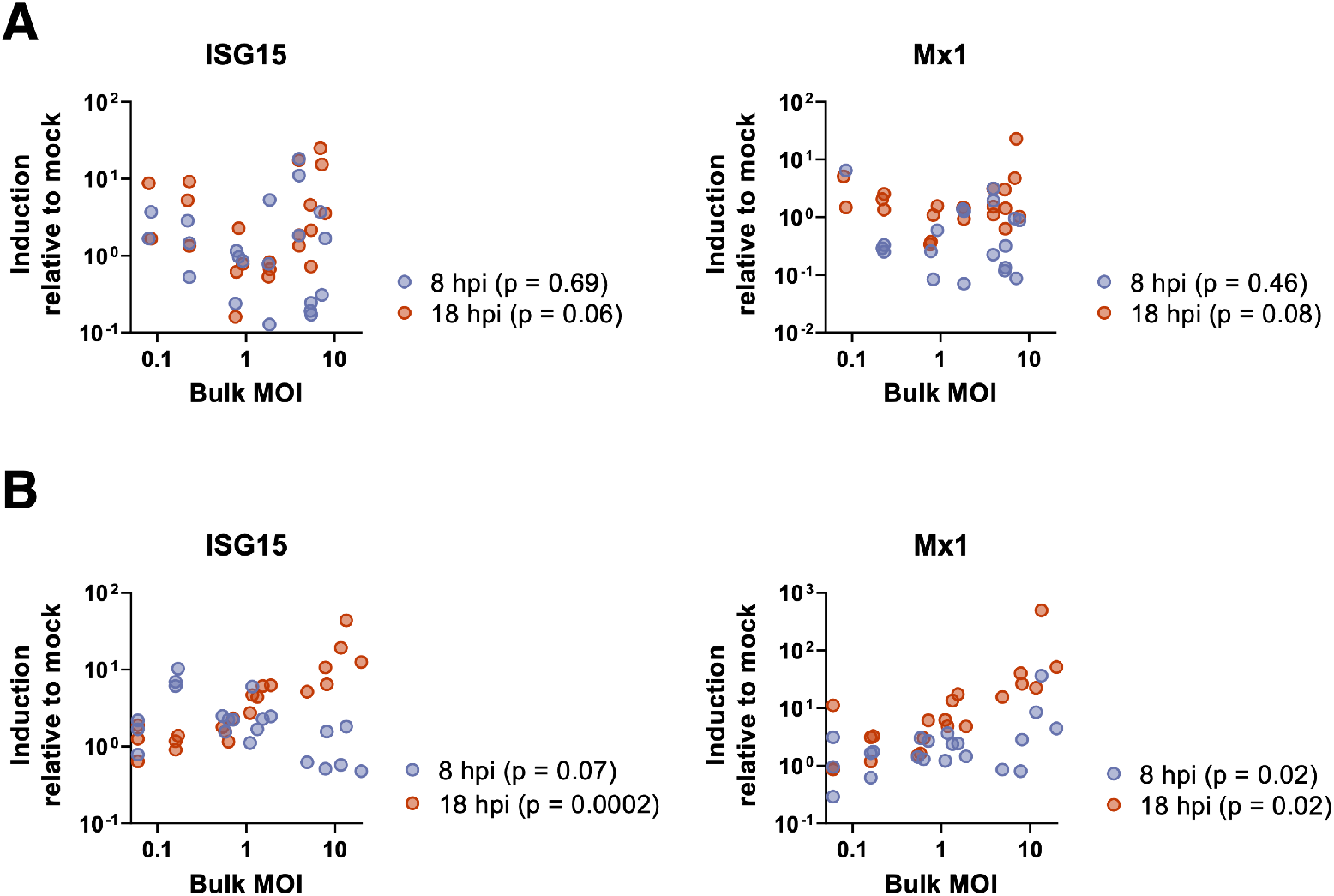
Cellular co-infection enhances ISG induction in A549 but not MDCK. **(A)** MDCK and **(B)** A549 cells were infected with PR8 under single cycle conditions at the indicated bulk MOIs. Levels of cellular ISG15 and M×1 transcript were measured by RT-qPCR at 8 and 18 hpi, compared to levels in mock cells. P values for the correlations between bulk MOI and fold induction of the indicated ISGs at the indicated timepoints are shown.

### Cellular co-infection enhances type III (but not type I) IFN induction in A549 cells

A prior study observed that the magnitude of interferon (IFN) transcription during IAV infection is higher under higher experimental MOI conditions (12); however, this relationship was not rigorously defined and it was not clear whether this effect simply arose from increases in infected cell numbers. In contrast, a more recent study suggests that increasing cellular MOI might result in more potent antagonism of IFN activation (15). To precisely define the effects of cellular co-infection on IFN induction, we measured the effects of varying bulk MOI on the induction of both type I (IFNB1) and type III (IFNL1) IFN transcription in both MDCK and A549 cells at 8 and 18 hpi under single cycle infection conditions.

For MDCK, expression levels of IFNB1 and IFNL1 in infected cells were barely if at all elevated, compared with mock cells (**Fig 7A**). We found no significant correlation between bulk MOI and levels of IFNB1 induction at either time-point tested (log-log linear regression: p = 0.38 for 8 hpi and p = 0.23 for 18 hpi). For IFNL1, expression was independent of bulk MOI (p = 0.67) at 8 hpi; however, at 18 hpi there was a statistically significant yet modest positive correlation between bulk MOI and levels of IFNL1 induction (slope = 0.34; p = 0.0077).

**Figure 7.**
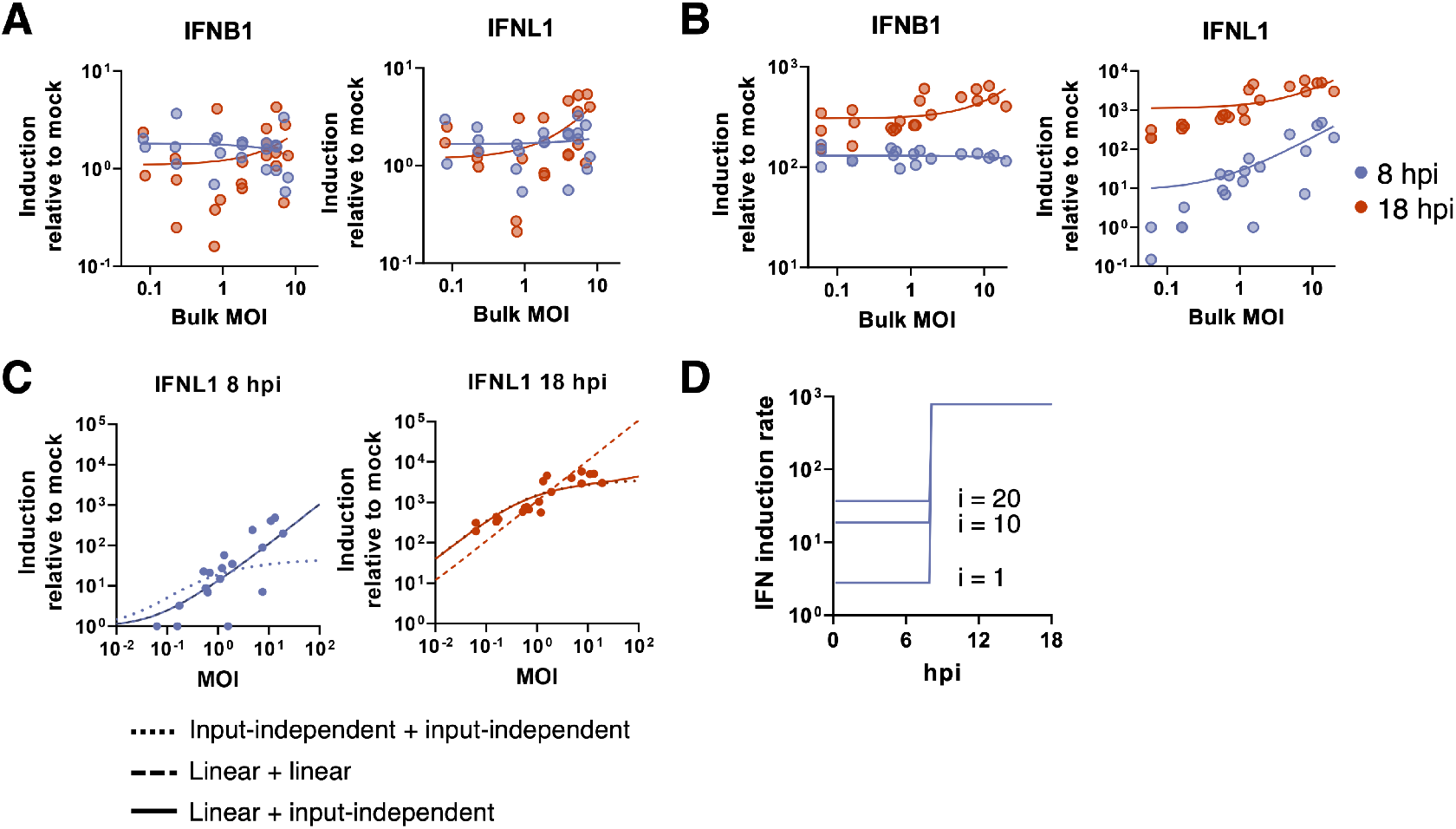
Cellular co-infection enhances type III (but not type I) IFN induction in A549 cells but has no significant effects on MDCK cell IFN induction. **(A)** Levels of cellular IFNB1 and IFNL1 transcript were measured by RT-qPCR at the times indicated in MDCK cells infected with PR8 under single cycle conditions at a range of bulk MOIs. Each data point represents values from a single infection well and lines represent log-log linear regressions. No significant positive correlation between IFNB1 induction and bulk MOI at 8 hpi and 18 hpi (p = 0.21 and p=0.24, respectively), and no significant positive correlation between IFNL1 induction and bulk MOI at 8 hpi and 18 hpi (p = 0.98 and p = 0.15, respectively). **(B)** Same as in (A) but for A549 cells. No significant correlation between IFNB1 induction and bulk MOI at 8 hpi (p = 0.81), but at 18hpi the slope is significantly nonzero (p =0.0002). For IFNL1, there is a significant positive correlation between normalized IFNL1 induction and bulk MOI at both timepoints (p < 0.0001). **(C)** Three IFN induction model fits to IFNL1 induction data in A549 cells at 8 and 18 hpi. Note that for the 0-8 hr plot, linear+linear and linear+input-independent are overlapping since they are the same for this epoch, and for the 8-18 hr plot, input-independent+input-independent and linear+input-independent are nearly overlapping. **(D)** IFN induction rates for the linear + input-independent model. From 0-8 hpi, IFN induction rates increase linearly with viral input: i = 1, 10, 20, and from 8-18 hpi, the IFN induction rate is independent of viral input.

In A549 cells, we observed that IFNB1 and IFNL1 expression responded very differently to increasing bulk MOI. By 8 hpi, IFNB1 expression was significantly elevated above mock under all MOI conditions (**Fig 7B**); however, there was no correlation between bulk MOI and levels of IFNB1 induction (p = 0.81). This suggests that IFNB1 induction, at least in bulk, is independent of MOI or even the total number of infected cells in the culture. By 18 hpi, we did observe a statistically significant positive correlation between bulk MOI and IFNB1 induction (p = 0.0002); however, this effect was quite small (slope = 0.16). In contrast to IFNB1 expression, we observed a strong positive correlation between IFNL1 expression and bulk MOI at both 8 and 18 hpi (p < 0.0001 for both time points; **Fig 7B**), although the effect was less pronounced at 18 hpi (slope = 0.58) compared with 8 hpi (slope = 1.08).

To determine whether the observed relationship between bulk MOI and IFNL1 induction in A549 cells could be explained simply as the result of increasing the number of infected cells, we statistically fit several mathematical models to these experimental data. Because only 8 hpi and 18 hpi data were available for model fitting, we considered different model combinations by piecing together a model governing the 0-8 hpi epoch with a model governing the 8-18 hpi epoch. During each epoch, we considered three different possible time-independent interferon induction rate models (**Table S5**; Methods): (1) an input-independent model in which all infected cells have similar IFNL1 induction rates relative to mock cells; (2) an input-dependent model in which IFNL1 induction rates scale linearly with virus input; and (3) an input-dependent model in which IFNL1 induction rates increase with virus input, saturating at high input values. The results of fitting these models jointly to the 8 hpi and 18 hpi IFNL1 induction data are shown in **Table S6**.

The best fitting model had IFNL1 expression in A549 cells be linearly dependent on cellular input during the 0-8 hr epoch, while being independent of cellular input during the 8-18 hr epoch. **Figure 7C** shows this combined model fit to the data, along with two other model combinations for comparison. A model assuming input-independence during both epochs does poorly in reproducing the 8 hpi data; in fact, all models with input-independence during the 0-8 hpi epoch do poorly (**Table S6**). A model that instead assumes linear input-dependence during both epochs can reproduce the 8 hpi data but does poorly at reproducing the 18 hpi data (**Fig 7C**). In **Fig 7D** we show graphically what the most supported IFN induction model looks like, for IFNL1 induction in A549 cells. Early in the infection (<8 hpi), the rate of IFNL1 induction depends linearly on cellular MOI, such that cells with higher levels of virus input have higher rates of IFNL1 induction than cells with lower levels of virus input. However, later in the infection (>8 hpi), the rate of IFNL1 induction does not depend on virus input. These results are consistent with a mechanism by which increasing cellular MOI accelerates the intra-cellular accumulation of viral replication products, resulting in earlier, more robust detection by RIG-I or other innate sensing pathways and a higher bulk rate of IFNL1 induction early during infection.

### Cellular co-infection decreases the potential for superinfection

Finally, we wanted to understand how cellular co-infection may affect the potential for cellular superinfection. We previously showed that under low bulk MOI conditions in multiple cells lines, increasing the number of functional gene segments delivered to a given cell resulted in more potent superinfection exclusion (SIE), limiting the potential for cellular co-infection in a cell-type independent manner (21). We thus hypothesized that increases in cellular MOI would shorten the time window during which superinfection is possible, thus limiting the total number of virions that can successfully infect a given cell.

To define how cellular MOI affects subsequent superinfection potential, we measured the extent of SIE across a range of bulk MOIs in MDCK cells. We generated 2 antigenically distinct reassortant viruses (rH3N1 and rH1N2) that could be differentiated using specific monoclonal antibodies. We infected MDCK cells with rH3N1 at a range of intended MOIs resulting in groups with average bulk MOIs of 0.25, 1.13, 3.00, 15.43, and 53.72 TCID50/cell (based on subtracting post-adsorption inoculum GE titers from pre-adsorption inoculum titers as detailed above). At 6 hpi, we superinfected with a constant bulk input MOI of rH1N2. To measure the baseline co-infection rates in the absence of SIE effects, we included co-infection controls for each input MOI where we simultaneously co-infected with rH3N1 and rH1N2 viruses. At 19 hpi, we examined the infection status of cells by flow cytometry, using H3 and H1 expression as markers of rH3N1 infection and rH1N2 infection, respectively.

As expected, during simultaneous co-infection (where no SIE occurs), we observed that the fraction of co-infected cells (H3+H1+) increased with the input MOI of rH3N1 virus and plateaued when almost all rH1N2-infected cells (H1+) were co-infected with rH3N1 (**Fig 8A**). These simultaneous co-infection data allowed us to develop and parameterize an appropriate “null” model of co-infection in the absence of SIE (**Methods**). To capture the possibility of overdispersion of viral particles (which we found to be important when analyzing cell death rates), the null model assumed that cells could belong to either a low susceptibility class of cells or a high susceptibility class of cells. **Fig 8A** shows the fit of this null model to the data.

**Figure 8.**
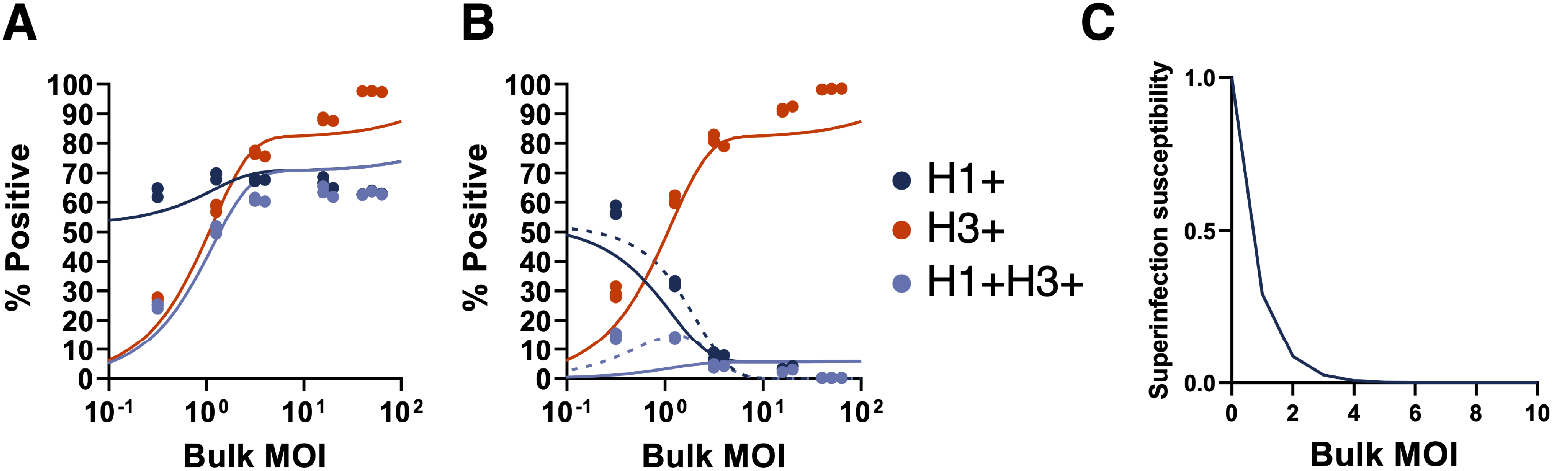
Cellular co-infection decreases the potential for superinfection. **(A)** MDCK cells were simultaneously co-infected with rH1N2 (at a constant MOI) and rH3N1 (at varying input MOIs shown; x-axis) under single cycle conditions; SIE would not be expected to occur during simultaneous co-infection. Plot shows the percentages of all cells infected with rH1N2 (H1+; includes co-infected cells; dark blue), all cells infected with rH3N1 (H3+; includes co-infected cells; red), and cells co-infected with both (H3+H1+; light blue), as determined by flow cytometry at 19 hpi. Solid lines indicate the two-susceptibility state null model fit to these data. **(B)** MDCK cells were infected with rH3N1 at varying input MOIs (x-axis) under single cycle conditions. 6 hours later, cells were superinfected with rH1N2 at an intended MOI of 0.5 TCID50/cell. Percentages of cells that were H1 + (including cells co-infected with rH3N1), H3+ (including cells co-infected with rH1N2), and H3+H1+ were determined by flow cytometry at 19 hpi. Lines indicate statistical fits of the input-independent (solid) and input-dependent (dashed) models. **(C)** Visualization of the input-dependent model of superinfection susceptibility where the susceptibility of infected cells to superinfection is shown relative to the susceptibility of uninfected cells.

In contrast to what was observed under simultaneous co-infection conditions, under superinfection conditions we observed that the fractions of both double-infected (H3+H1+) cells and rH1N2-infected (H1+) cells decreased with increasing rH3N1 input MOI. To determine whether these patterns could simply be explained by higher numbers of cells being infected with rH3N1 at higher rH3N1 input MOIs, we assessed how well two different models of super-infection fit the experimental data (Methods). The first model assumed that all rH3N1-infected cells had the same reduced probability of becoming infected with rH1N2 (input-independent). The second model assumed that the probability of being infected with rH1N2 decreased with cellular rH3N1 MOI (input-dependent). Similar to the co-infection null model, both models assumed that cells fall into either a high or low susceptibility class. We fit each of these two models to the SIE data. The input-independent model underestimated the extent of rH1N2 infection to a greater extent than the input-dependent model (**Fig 8B**). The input-independent model further systematically underestimated the extent of double (H3+H1+) infection at low rH3N1 input MOI, while systematically overestimating these measurements at high rH3N1 input MOI (**Fig 8B**). In contrast, the input-dependent model was better able to reproduce the double (H3+H1+) infection measurements across all experimental MOIs (**Fig 8B**). As anticipated from these patterns, we found that the input-dependent model was significantly better supported by the data than the input-independent model (**Table S7**). **Fig 8C** shows the relationship between cellular input and susceptibility to superinfection predicted by this input-dependent model, indicating a rapid decline in susceptibility to superinfection with rH1N2 at 6 hpi at increasing levels of rH3N1 virus input.

These results are consistent with our previous finding that susceptibility to superinfection is inversely correlated with the cellular dosage of replication complexes delivered by incoming virions (21). Thus, the MOI-dependence of SIE may serve as a negative feedback loop that restricts the maximum number of virions that can successfully infect a given cell.

## DISCUSSION

In this study, we defined the functional consequences of cellular co-infection by quantifying the effects of cellular MOI on the phenotypes of infected cells. We combined precise, quantitative single-cycle *in vitro* infection experiments with statistical model fitting to demonstrate that at the cellular level, variation in viral input gives rise to substantial variation in infection outcomes. This includes cell line-dependent variation in viral output dynamics as well as the host transcriptional response to infection. Intriguingly, type I and type III IFN exhibited distinct responses to increases in MOI, suggesting that variation in cellular MOI could alter the balance of these two antiviral cytokines. Altogether, these results clearly suggest that the incidence of cellular co-infection could play an important role in influencing influenza virus infection outcome.

These results complicate the common understanding of viral genotype-phenotype relationships. Generally, we understand that the phenotype of a virus within a given host system is encoded by its genome sequence. Our data add another layer to this view: it is not simply the sequence(s) of a viral genome that influences its phenotype but also how those sequences are distributed across cells. Two viruses with identical genome sequences could exhibit significantly different replication dynamics or patterns of IFN induction if they differ in their cellular MOI distribution (22,23). These results suggest that we need to better understand the forces that govern the extent of cellular co-infection and the spatial distribution of virions during IAV infection.

Our finding that the relationship between cellular MOI and viral replication efficiency differed between cell types complements the results of two recent reports. In the first, Andreu-Moreno et al. examined the effect of cellular co-infection on vesicular stomatitis virus (VSV) fitness across multiple cell types (22). They found that increasing cellular MOI (via virion aggregation) enhanced viral fitness, but that the magnitude of this effect varied between cell types. Specifically, they observed an inverse relationship between the susceptibility of a given cell line to VSV infection and the extent to which cellular co-infection enhanced viral output. Similarly, Phipps et al. observed that cellular co-infection could enhance influenza virus replication when the virus strain used was poorly adapted to the cell line used (20). Along with these studies, our results illustrate the importance of considering variation between cell types. This point is especially relevant given the diversity of cell types infected by IAV *in vivo*. Beyond cell type variation, it is likely that other viral genotypes will exhibit quantitative or qualitative differences in the dynamics that we have measured here. Thus, we do not argue that our specific results will describe all combinations of influenza virus strains and target cell types. Instead, our results identify cellular co-infection as an important determinant of viral replication dynamics and immune activation and provide a rigorous quantitative framework for future efforts to understand the consequences of co-infection.

It is important to point out that the experimental data described here were all collected from aggregate populations of cells and that our models ignore single cell heterogeneity between cells with the same cellular MOI. There is substantial heterogeneity in viral gene expression and progeny production between individual cells infected under similar conditions (24–27). Time-course experiments from single poliovirus-infected cells have demonstrated that bulk measurements like those used here can obscure more complex, heterogeneous dynamics that occur at the single cell level (28). A more complete understanding of the functional consequences of cellular co-infection will likely depend upon future studies that quantify single cell variability.

The IFN system represents one of the earliest and most potent lines of defense against IAV within the respiratory tract (29–31). Similar to previous reports, we observed that IAV infection resulted in a more dramatic induction of type III IFN (32), compared with type I IFN, in A549 cells. We were surprised to find that type I and type III IFN induction responded differently to increases in input MOI, as the induction pathways that lead to initiation of IFNB1 and IFNL1 are thought to largely overlap (33). Our data suggest that the induction or regulatory circuitry for these two pathways are differentially affected by the amount of viral input. Even more surprising was our finding that type I IFN induction was largely insensitive to input MOI over two orders of magnitude. In our hands, not only was IFNB1 induction MOI-independent at the cellular level, it was also largely unaffected by the total number of infected cells, which ranged from ~3% to ~63% across the samples tested. Induction of IFNB1 expression by IAV is known to be highly stochastic at the single cell level and many infected cells do not upregulate IFN expression (15,24,34); however, it is still counterintuitive that increasing the total number of infected cells would not increase the overall number of cell producing IFNB1 (and thus the bulk expression level). More work is clearly needed to understand the factors that govern IFN expression at the single cell level.

The implications of these results for understanding what happens during natural infection remain unclear, but at a minimum suggest that frequent occurrence of cellular co-infection is likely to boost the magnitude of the type III, but not the type I IFN response. This dynamic could lead to MOI-dependent changes in the relative balance of type I versus type III IFN induction which could be consequential given that these cytokine families can trigger non-redundant effector responses during IAV infection (35–38). One important caveat here is that we are only examining IFN induction within a single cycle of replication and a single cell type and patterns of type I and type III IFN induction over the course of infection are sure to be much more complicated.

Given that cellular MOI and co-infection have clear functional consequences during IAV infection, it is critical to quantify the actual occurrence of cellular co-infection during *in vivo* infection and to identify host and viral determinants of cellular co-infection frequency. Natural infections are likely typically initiated under low MOI conditions due to the relatively small size of transmitted founder populations (though even small numbers of transmitted virions could achieve high cellular MOI if they are physically aggregated) (39,40). Even if infection is initiated at low cellular MOI, high cellular MOI conditions could be very rapidly established if viral spread is locally restricted, as appears to be the case (8). This idea is supported by a recent study that demonstrated a virus that is entirely dependent upon cellular co-infection to replicate can successfully replicate in guinea pigs (7). Though very few studies have attempted to actually quantify the MOI distribution *in vivo* because of the inherent technical challenges, our work suggests that this feature could be an important, underappreciated determinant of infection outcome.

This study also highlights the power of interfacing carefully quantified experimental data with statistical modeling approaches. By fitting models to our experimental data, we were able to rigorously test our hypotheses concerning the phenotypic effects of cellular MOI to an extent that would not be possible based on standard data analysis. This was essential for confidently distinguishing between the different functional forms that could potentially describe the phenotypic consequences of different levels of virus input. Hopefully, the data generated here will help inform future mathematical modeling efforts focused on understanding IAV within-host infection dynamics. Our results to date certainly suggest that future model structures may better capture the underlying biology of IAV infection if they account for the phenotypic consequences of cellular co-infection (41,42).

Altogether, our results clearly demonstrate that cellular MOI can have concrete effects on infection outcome, highlighting the functional importance of collective interactions during IAV infection (43). At a minimum, our data establish that the distribution of viral genomes across cells and the patterns of cellular co-infection that result can significantly alter emergent viral infection phenotypes. This suggests that future efforts to understand influenza virus infection dynamics and outcomes should consider patterns of virus input and cellular co-infection.

## MATERIALS AND METHODS

### Cells

HEK293T (human embryonic kidney) cells were used for rescue of PR8 by plasmid transfection. MDCK (Madin-Darby canine kidney) cells were used for virus propagation, TCID_50_, and infection experiments. 293T and MDCK were maintained in MEM (Thermo Fisher Scientific) supplemented with 8.3% fetal bovine serum (FBS). A549 (human lung carcinoma) cells were used for infection experiments and were maintained in Ham’s F12 nutrient mixture (Thermo Fisher Scientific) with 8.3% FBS.

### Virus rescue and propagation

Eight plasmids encoding the A/Puerto Rico/8/1934 (H1N1; PR8) segments PB2, PB1, PA, HA, NP, NA, MA, and NS in the dual promoter pDZ vector were used to generate recombinant virus stocks through standard IAV reverse genetics. The recombinant PR8 virus (rPR8) differs from the published sequence (GenBank accession nos. AF389115–AF389122) at two positions: PB1 A549C (K175N) and HA A651C (I207L) (numbering from initiating Met). rH3N1 and rH1N2 viruses used in superinfection studies were similarly generated through reverse genetics, using the HA and NA segments from A/Udorn/307/72 (H3N2) respectively and the remaining segments from PR8.

For virus rescue, 500 ng of each plasmid were mixed with 200 *μ*L jetPRIME buffer (Polyplus-transfection) and 8 *μ*L jetPrime reagent and incubated at room temperature for 10 minutes. The transfection mixture was added dropwise to 293T cells at 60% confluency in 6-well cell culture plates and incubated at 37°C, 5% CO2. Media was changed 18-24 hours post-transfection, with 2 mL of virus growth media (MEM, 1 mM HEPES, 1 *μ*g/mL TPCK trypsin (Worthington Biochemical Corporation; Lakewood, NJ, USA), 50 *μ*g/mL gentamicin) containing 2.5×10^5^ MDCK cells. Rescue supernatant was harvested and used in plaque assay. Briefly, 300 *μ*l 10-fold serially diluted transfection supernatant in 1X DPBS (+Ca/+Mg), 0.1% BSA, pH 7.1, was added per 100% confluent MDCK well in 6-well cell culture plate, in duplicate. After 1 hour of incubation, 3 mL agarose overlay (0.9% agarose, 1X EMEM, 1 *μ*g/mL TPCK trypsin, 0.2% BSA, 1X glutamax, 1 mM HEPES, 50 *μ*g/mL gentamicin) was added to each well and incubated for 48-72 hours at 37°C.

To generate seed virus stocks, three isolated plaques were picked and resuspended in 200*μ*L 1X DPBS (+Ca/+Mg), 0.1% BSA; 100 *μ*L was added to one well of 6-well cell culture plate with 80% confluent MDCK cells with 2 mL of virus growth media and incubated at 37°C, 5% CO2 for 48-72 hours. To generate working stock virus, confluent MDCK cells were infected with seed virus stock at an MOI of 0.001 TCID_50_. One-hour post infection, inoculum was discarded and replaced with virus growth media and incubated at 37°C, 5% CO2 for 48-72 hours. Supernatant was clarified at 3000 RPM for 10 minutes and 500 *μ*L aliquots were stored at −70°C.

50% tissue culture infective dose (TCID_50_) for PR8 viral supernatant was titrated in 96-well cell culture plates with 100% confluent MDCK cells by 1:10 serial dilution in virus growth media. Cytopathic effect was visualized 3-5 days post infection. TCID_50_ values were calculated based on Reed and Munch calculation (44).

### Viral output assay

To precisely quantify the effects of viral input on viral output and cell death rates, PR8 was diluted in 1X PBS +/+ pH 7.3 in 200 *μ*L triplicates to yield intended MOIs of 0.25, 1,2.5, 5, and 25 and used to inoculate MDCK or A549 cells in 24-well plates. Infections were synchronized by carrying out adsorption at 4°C for 1 hour. The supernatant was collected at 1 hpi, the cell monolayers were washed three times with 1X PBS, media was replaced with 500 *μ*L MEM or F12, for MDCK and A549 respectively, with 8.3% FBS and cells were shifted to 37°C. At 2 hpi, media was supplemented with 20 mM NH_4_Cl to block secondary spread of the virus (45). Total viral supernatant was collected at 3, 6, 12, and 18 hpi and stored for at −70°C for later use. Cell monolayers were then trypsinized and either stained with trypan blue and manually counted with a bright-line hemocytometer (to quantify remaining live cells) or prepared directly for flow cytometry.

### Quantification of infected cell percentages

To quantify infected cells, cells in suspension were transferred to 96-well plate round bottom plate, centrifuged for 5 minutes at 1000 RPM, and the supernatant was removed by one quick flick. Cells were washed once with 200 *μ*L 1X PBS with 0.1% BSA, then 200 *μ*L 1X FoxP3 transcription factor fixation/permeabilization (Thermo Fisher Scientific) was added to each well and incubated on ice for 30 minutes or 4°C overnight while protected from light. Cells were then washed three times with perm wash, 1X PBS with 0.1% BSA and 0.1% saponin. 100 *μ*L primary monoclonal antibodies anti-HA (H28-E23-AF488) and anti-NP (HB65-AF647) were incubated with each sample for 30 minutes on ice, washed three additional times to remove unbound antibody, and resuspended in 200 *μ*L perm wash. Labeled cells were immediately analyzed with BD LSR II flow cytometer (BD Biosciences) and FlowJo version 10.4 software package (FlowJo, LLC). Positively infected cells are any cells that expresses one or both protein of interest (HA or NP); mock infected cells were used to gate for negatively infected cells. The negative gate for MDCK cells at MOI 5.35, 5.41, and 5.45 was shifted to account for the shift in fluorescence seen in uninfected cells.

### Measurement of viral genome equivalents by RT-qPCR

Viral RNA was isolated from viral supernatant with Zymo vRNA 96-well extraction kit (Zymo Research) according to manufacturer’s instructions, eluted with 30 *μ*L RNase free water, and stored at −70°C. For cDNA synthesis, 5 *μ*L vRNA, 0.5 *μ*L 10 mM dNTP mix (Sigma-Aldrich), and 1.0 *μ*L Uni12(46) (AGCAAAAGCAGG) were incubated at 65°C for 5 minutes then transferred to ice for 2 minutes. 1 *μ*L SUPERase In RNase inhibitor (20 U/*μ*L; Thermo Fisher Scientific) was added to each mixture and incubated on ice again for 2 minutes. 6.5 *μ*L dH_2_O, 4 *μ*L 5X First-Strand Buffer, 1 *μ*L 100 mM DTT, and 1 *μ*L SuperScript III RT (200 U/*μ*L; Thermo Fisher Scientific) were added to each reaction and incubated at 55°C for 60 minutes and heat inactivated at 70°C for 15 minutes.

Genome equivalents (GE) were estimated by RT-qPCR of MA gene segment. In duplicate for each sample, 10 *μ*L Power SYBR Green PCR master mix (Thermo Fisher Scientific), 0.5 *μ*L 10 *μ*M forward and reverse primers (MA-Forward: ACAGAGACTTGAAGATGTC and MA-Reverse: TCTTTAGCCATTCCATGAG), 8 *μ*L dH_2_O, and 1 *μ*L cDNA were added to 0.2 mL MicroAmp Optical 96-well reaction plate (Thermo Fisher Scientific). RT-qPCR was performed on the QuantStudio 3 (Thermo Fisher Scientific) platform and the cycling conditions were as follows: 95°C for 10 minutes, 40 cycles of 95°C for 15 seconds and 60°C for 60 seconds. The standard curve established with pDZ-PR8 MA plasmid (R^2^ of 0.9992) was used to estimate GE/*μ*L and then each sample was corrected for dilution factor to give a final GE/mL.

### Superinfection assay

For the 6hr superinfection group, confluent MDCK cells in six-well plates were infected with rH3N1 virus at intended MOIs of 0.05, 0.25, 1,2.5, and 10 TCID50/cell respectively at 4°C. 1 hour post-adsorption, remaining inoculum was collected, monolayers were washed with PBS and incubated in MEM + 8.3% FBS. At 6 hpi, monolayers were superinfected with rH1N2 at MOI=0.5 TCID50/cell at 4°C. One hour post-adsorption, monolayers were washed with PBS and incubated in MEM + 8.3% FBS. At 9 hpi of rH3N1 (3 hpi of rH1N2), the media was changed to MEM with 50 mM HEPES and 20 mM NH4Cl to block spread of both viruses. For the 0hr co-infection group, cells were infected with a mixture of rH3N1 and rH1N2 at the same MOIs as in 6hr superinfection group. At 3 hpi, 20 mM NH4Cl was added to block viral spread. For both 0hr and 6hr groups, at 19 hpi of rH3N1 (13 hpi of rH1N2), cell monolayers were trypsinized into single-cell suspensions and stained with Alexa Fluor 647-conjugated mouse anti-N1 mAb NA2-1C1 and Alexa Fluor 488-conjugated mouse anti-H1 mAb H28-E23 (gifts of Dr. Jon Yewdell). After staining, cells were washed with PBS, run on a BD LSR II, and analyzed using FlowJo version 10.1 (Tree Star, Inc.). To quantify actual MOIs, virus present in both pre- and post-adsorption inoculum was quantified by RT-qPCR as above using the following primers specific for the N1 segment: AAATCAGAAAATAACAACCATTGGA, ATTCCCTATTTGCAATATTAGGCT.

### Interferon quantification and interferon stimulated gene quantification

Interferon β1 (IFNB1) and interferon λ1 (IFNL1) and interferon stimulated genes (ISG15 and Mx1) induction was quantified during infection of MDCK and A549 cells with PR8 at intended MOIs of 0.1-100. In brief, MDCK and A549 cells in 24-well plate were inoculated for one-round of replication, as described above. At 8 or 18 hpi, supernatant was removed, cells were washed thoroughly, dissociated from the plate, and cellular mRNA was extracted with RNeasy Mini kit (QIAGEN), according to manufacturer’s instructions. Reverse transcription was completed with SuperScript III first-strand synthesis system for RTq-PCR (Thermo Fisher Scientific). Briefly, 8 *μ*L cellular mRNA was incubated with 2.5 ng/*μ*L random hexamers, and 0.5 nM dNTP mix at 65°C for 5 minutes then placed on ice for 1 minute. Then 1X RT buffer, 5 mM MgCl2, 0.01 M DTT, 2 U/*μ*L RNaseOUT, and 10 U/*μ*L SuperScript III RT were added to each sample and incubated at 25°C for 10 minutes, 50°C for 50 minutes, and 85°C for 5 minutes; finally 1 *μ*L RNase H was added and incubated at 37°C for 20 minutes. With cellular cDNA for A549, TaqMan gene expression assays (FAM-MGB; Thermo Fisher Scientific) specific for IFNB1 and IFNL1 were used in multiplex with TaqMan GAPDH control reagents kit (JOE-TAMRA; Thermo Fisher Scientific). Briefly, in duplicate 2*μ*L cellular cDNA was added to 2 *μ*L TaqMan Fast Advanced Master Mix, 1 *μ*L either IFNB1 or IFNL1 TaqMan Gene expression assay, 100 nM GAPDH forward and reverse primers, 200 nM GAPDH TaqMan probe, and dH_2_O in 20 *μ*L. RTq-PCR was performed on the QuantStudio 3 platform and the cycling conditions were as follows: 95°C for 20 seconds, 40 cycles of 95°C for 1 second and 60°C for 20 seconds. For the remaining targets of both MDCK and A549, primers were used for SYBR green chemistry (see **Table S8**), with β-actin as an endogenous control. Briefly, in duplicate 2 *μ*L cellular cDNA was added to 10 *μ*L PowerUp SYBR Green Master Mix (Thermo Fisher Scientific), 250 nM of each forward and reverse primers, and dH_2_O in 20 *μ*L. RTq-PCR was performed on the QuantStudio 3 platform and the cycling conditions were as follows: 95°C for 10 minutes, 40 cycles of 95°C for 15 seconds and 60°C for 1 minute, followed by melt curve analyses of 95°C for 15 seconds, 60°C for 1 minute and 95°C for 1 second. Induction of IFN and ISG was quantified by the fold change of infected cells from mock infected cells.

### Statistical modeling of cell death

To infer infected cell death rates from experimental data, and to determine whether these rates are time-dependent and/or virus input-dependent, we fit the following general model to the experimental data points from 3, 6, 12, and 18 hpi:

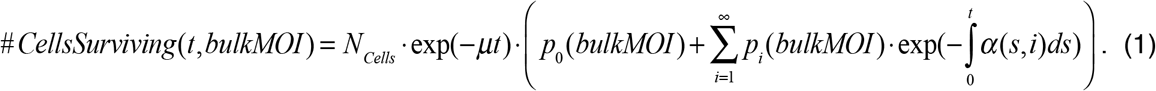

In this equation, *N*_cells_ denotes the initial number of cells on the plate, the parameter *μ* denotes the background death rate of (both infected and uninfected) cells, and *α*(*s,i*) is the general form for the instantaneous death rate of of cells at time *s* since infection, infected with viral input *i*.

*p_i_*(*bulkMOI*) denotes the initial fraction of the cell population that has viral input *i*, which depends on the actual bulk MOI and the model specifying the distribution of viruses across cells. We included a background cell death rate *μ* because we noted a small (but non-negligible) amount of cell loss in mock infected (uninfected) cells. Because of uncertainty in the initial number of cells, we estimate *N_cells_*. along with the other parameters in the model fitting to experimental data.

This expression assumes that uninfected cells (*i* = 0) survive the time course of the experiment with probability exp(−*μt*). The probability of infected cells with viral input *i* surviving up to time *t* is given by the survival function, 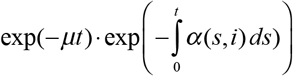. The percent of surviving cells that are infected at time *t* is given by

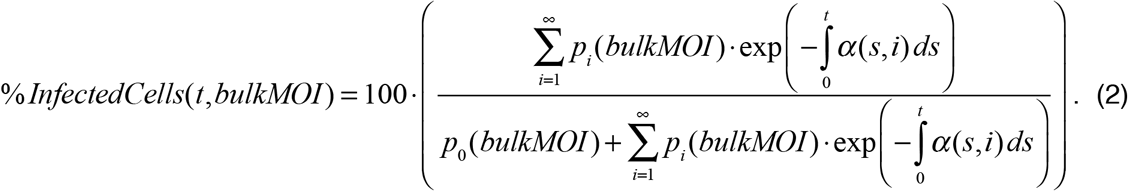

For each of the four cell death rate models listed in **Tables S1** and **S2**, we estimated the parameter *μ* and the parameters of *α*(*s,i*) by simultaneously fitting models to the time course data on the number of cells surviving and to the flow data on the percent of infected cells at *t* = 18 hpi. We performed these fits each three times, under three different viral distribution models: a Poisson distribution, a negative binomial distribution, and a zero-inflated Poisson distribution. For both the Poisson and negative binomial distributions, we parameterized models with the mean of the distribution equaling the mean actual bulk MOI. When fitting to the data under the assumption of a negative binomial distribution, we further simultaneously estimated *r* along with *μ* and the parameters of *α*(*s,i*). At *r* = ∞, the negative binomial distribution is equivalent to the Poisson distribution. As *r* becomes smaller (closer to 0), the distribution of virions become increasingly overdispersed. For the zero-inflated Poisson distribution, the mean of the zero-inflated Poisson distribution is (1 − *p*) · *bulkMOI*, where *p* is an estimated parameter between 0 and 1 that quantifies the probability of extra zero counts. When fitting the zero-inflated Poisson model, we thus estimated *p* along with m and the parameters of *α*(*s,i*). All viral infectivity distributions considered (Poisson, negative binomial, and zero-inflated Poisson) fed into the evaluation of *p_i_*(*bulkMOI*).

The first model of cell death rate we considered was a time-independent, input-independent model, such that cell death rates were the same across all infected cells and cell death rates did not change over the course of a cell’s infection. The second model we considered was a time-dependent, input-independent model. For this model, we let the infected cell death rate be given by the Weibull hazard function, a commonly used function to incorporate time-dependency into time of death estimations. The Weibull hazard function is a two-parameter model (*b, k*) with functional form given in **Table S1**. An estimate of the shape parameter *k* of less than 1 indicates that the death rate decreases over time, while an estimate of *k* greater than 1 yields a death rate that increases over time. When *k* = 1, this time-dependent, input-independent model becomes identical to the time-independent, input-independent.

The third and fourth models we considered allowed for cell death rates to depend on viral input: the first of these two models assumed that cell death rates were time-independent; the second further allowed for time-dependency, again according to the Weibull hazard function (**Table S1**). The input-dependency was incorporated into the third and fourth models using the monomial *i^ε^* where *i* is again viral input (*i* = 1,2,3,…) and *ε* > 0 is real-valued. The time-independent, input-dependent model of cell death rate has the form *c* · *i^ε^*, where *c* is a positive constant. When *ε* = 0, this model becomes identical to first time-independent, input-independent cell death rate model. The fourth model, the time-dependent, input-dependent model, has the form *α*(*s, i*) = *d* · *i^ε^* · *k* · *s*. When *ε* = 0, the model becomes identical to the time-dependent, input-independent model, i.e. the second model considered. When *ε* = 0 and *k* = 1, the model becomes identical to the time-independent, input-independent model, i.e. the first model considered. When *ε* > 0 and *k* = 1, this model is identical to the time-independent, input-dependent model, i.e the third model.

### Statistical modeling of virus production

To infer rates of virus production from infected cells and to determine whether these rates depend on cellular input or time since cellular infection, we developed six virus production models (**Table S3**) and statistically fit these models to the data shown in **Figure 4**. When fitting each of the virus production models to these data, we incorporated parameter estimates of cell death rates and viral infection distributions that were most supported by the data, in a cell line-specific manner. Specifically, for both MDCK and A549 cells, we let viral input be distributed across cells according to a negative binomial distribution and let the cell death rate be given by a Weibull hazard function (parameters provided in **Table S1,S2**).

We considered a set of six virus production rate models, *ν*(*t, i*), which are listed in **Table S3**. The general equation governing the total amount of virus produced in bulk cell culture at *t* hours post infection is given by

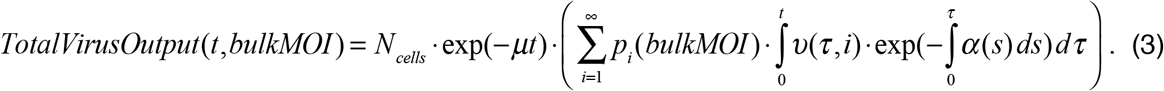

Here, the parameter, *N_cells_*, is the initial number of cell which was based on quantification of the total number of cells used in the bulk cell culture experiments of virus output, which was 2 million cells.

### Statistical modeling of the interferon response

To infer whether the rate of IFN induction from a cell was dependent on cellular MOI, we fit models of the interferon response (**Table S5**) to the IFN induction levels (relative to mock) that are shown in **Figure 7**. Because there was not a clear dependence of IFNB1 induction on bulk MOI and because there was no appreciable IFN induction in MDCK cells, we decided to fit models only to the IFNL1 induction data from A549 cells. While fitting these IFN response models, we assumed, based on our earlier analyses, that virions were distributed according to a negative binomial distribution and that infected cells died according to the time-dependent, input-independent model of cell death rate (parameters provided in **Table S2**). Because only two time point measurements were available, we fit the IFN response data using a piecewise approach, rather than considering continuous time models of IFN induction. Specifically, we assumed that during each of the two epochs (0-8 hpi and 8-18 hpi), IFN induction occurred at a rate that was constant in time but that could depend on cellular MOI. The three different models we considered implemented different forms for MOI-dependency. Because each epoch could be governed by one of three models, a total of 9 models were fit to the data shown in **Figure 7C**.

The general model equation for the level of IFN induction (relative to mock) at 8 hpi is:

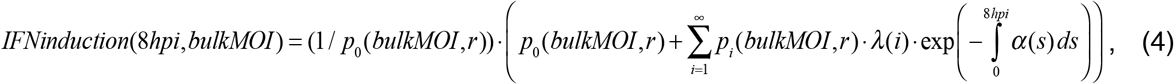

where *λ*(*i*) is the induction rate model as a function cellular MOI, *α*(*s*) is the time-dependent, input-independent model of cell death rate for infected A549 cells, and *r* is the dispersion parameter of the negative binomial distribution. The parameters associated with the time-dependent cell death rate and virus distribution models were set to the estimates given in Table S2.

The general model for IFN expression (relative to mock) between 8 and 18 hpi is given by:

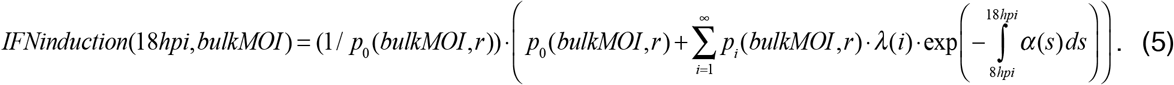

The level of IFN expression (relative to mock) predicted by the model is given by the model-predicted level of IFN expression by 8 hpi, plus the model-predicted level of IFN expression between 8 and 18 hpi.

We considered all nine possible model combinations (3 models for 0-8 hpi epoch x 3 models for 8-18 hpi epoch), fitting each simultaneously to INFL1 data at 8 and 18 hpi (**Table S6**). The best model combination was the model that assumed linear dependence of viral input on IFN induction during the 0-8 hpi epoch and cellular MOI-independence during the 8-18 hpi epoch. (**Table S6**; **Fig 7C,D**).

### Statistical modeling of superinfection exclusion

We first developed and parameterized an appropriate ‘null’ model, where both viruses are introduced simultaneously, and superinfection exclusion would not be anticipated. In this model, we assumed that cells differed in their susceptibility to viral infection. This assumption reflects our finding that viral particles are unlikely to be Poisson-distributed across cells (**Table S1**). In our null model, we did not adopt a negative binomial model to implement the possibility of viral particle overdispersion, as we had earlier, because if both reassortant viruses were assumed to be distributed according to a negative binomial distribution, viral particles (together) would not be distributed according to a negative binomial distribution. We instead considered a model that implemented different cell susceptibility classes, which we found to better accommodate viral overdispersion of distinct viral strains. To maintain simplicity, we considered only two types of cells: cells that had high susceptibility to infection and cells that had low susceptibility to infection. We defined fraction *y* of the cell population to be in low susceptibility state; the remaining fraction (1 - *y*) we assumed was in a high susceptibility state. We let a fraction *x* of the viral population enter the low susceptibility state cells; the remaining fraction of the viral population (1-*x*) we assumed entered the high susceptibility state cells. Under this model, the MOI of specifically the low susceptibility class of cells is given by (input MOI)*(*x*/*y*), and the MOI of specifically the high susceptibility class of cells is given by (input MOI)*(1 - *x*)/(1 - *y*). When *x* = *y*, the Poisson distribution assumption is recovered, and both classes of cells have the same input MOI. We fit this simple model to the co-infection data, estimating three parameters: the actual rH1N2 MOI, and the fractions *x* and *y*. The parameter values, estimated using an RSS approach, are actual rH1N2 MOI = 1.86 (95% CI = [1.71,2.03]), *x* = 2.14e-4 (95% CI = [0.334e-4, 13.8e-4]), and *y* = 0.0526 (95% CI = [0.0381,0.0725]). This parameterized model fit the experimental data well (**Fig 7A**). Since the parameter *x* was estimated to be close to 0, this model effectively implemented the zero-inflated Poisson model shown in Table S1, which had slightly lower statistical support than the negative binomial distribution.

To analyze the data from the superinfection experiment, we set as given the three parameter values and simple two-state susceptibility model structure derived from the fitting of the data from the simultaneous co-infection experiment described above. We then considered two distinct models to determine how cellular MOI may impact the rate of superinfection exclusion: an input-independent model and an input-dependent model. The input-independent model assumed that all infected cells had the same lower chance of being superinfected than previously uninfected cells. The parameter *s* quantified the extent of susceptibility of the previously infected cells (1 being full susceptibility). The input-dependent model instead assumed that cells that were infected with rH3N1 could experience different probabilities of superinfection exclusion. These different probabilities depended on a rH3N1 virus input, with, presumably, higher levels of rH3N1 virus input corresponding to higher probabilities of superinfection exclusion. For the input-dependent model, we specifically assumed a functional form given by *r^i^*, where *i* denotes rH3N1 virus input and 0 ≤ *r* ≤ 1. We estimated *s* for the input-independent model to be 0.0361 (95% CI = [0.0227, 0.0576]) and *r* for the input-dependent model to be 0.293 (95% CI = [0.214, 0.402]). Based on AIC, the input-dependent model is strongly preferred over the input-independent model (ΔAIC = 22.0) (**Table S7**).

### Fitting models to data and model selection criterion

To fit models to data, we minimized the sum squared error between the models and data to obtain the residual sum of squares (RSS). Note that minimizing the sum squared error is equivalent to maximizing the log-likelihood assuming normally distributed measurement error. These minimizations were done on the linear scale for the percent cells surviving and flow data, on the natural log scale for the virus output data, and on the log base 2 scale for IFN response to reflect the likely scale at which measurement error occurs for these data.

We calculated 95% confidence intervals by numerically estimating the covariance matrix using the “sandwich estimator.” This method approximates the observed Fisher information matrix by numerically estimating the inverse of the Hessian and allowing for the residuals to have different variances (47). We accounted for the chain rule in the log-transformation of parameters to obtain partial derivatives when estimating 95% percent confidence intervals. When parameter estimates are very large (e.g. the saturating input-dependent IFN induction models from 0-8 hpi; **Table S6**) or very close to zero (e.g. the input-dependent models of cell death under the negative binomial distribution; **Tables S1**, **S2**), the estimator of the covariance matrix is close to singular and thus sample variance estimates obtained from numerically inverting this matrix may be unreliable. In the more complex set of models, these large and close-to-zero estimates indicate further support for a simpler model that is nested within the more complex one.

To perform model selection, we used the Akaike Information Criterion (AIC). AIC based on RSS values is given by the equation: 2*k* + *n* ln(RSS) + constant, where *k* is the number of estimated parameters and *n* is the number of data points (48). Since AIC is a relative measure of information loss and the model with the lowest AIC has the most support, we calculated ΔAIC values to perform model selection by taking the difference between a given model and the model with the lowest AIC value.

In the tables throughout, we report the fitting results of each set of models, including parameter estimates with 95% confidence intervals, minimal sum squared error (RSS), and ΔAIC values.

All statistical modeling code is available on GitHub: https://github.com/Jeremy-D-Harris/MOIpaper_models_data.

## ACKNOWLEDGMENTS

Work in the Brooke and Koelle labs on this study was generously supported by DARPA INTERCEPT contract (W911NF-17-2-0034). The Brooke lab was further supported by an NIAID grant (1R01AI139246-01A1) on this study.

## SUPPLEMENTAL FIGURE LEGENDS

**Fig S1. FACS quantification of infected cell percentages based on HA and NP expression.** for MDCK (top row) and A549 (bottom row) cells. Gates for determining infection status were drawn based on NP and HA expression of mock cells. Gates were modified by eye for the MDCK cell line at the MOI of 5.35 for MDCK and 7.81 for A549 to better exclude negative cells. These data were generated from the same experiments used to generate cell death and virus production data.

**Figure S2. MDCK cell survival patterns cannot be reproduced under a time-independent, input-independent cell death rate model. (A)** The number of cells remaining for 3, 6, 12, and 18 hpi, respectively, as a function of bulk MOI, along with time-independent, input-independent cell death rate model fits (lines). **(B)** Number of surviving MDCK cells that are infected at 18 hpi, as measured by FACS, along with the negative binomial distribution model fit (line). As in Figure 2C, statistical parameterization of this model (overdispersion parameter *r* = 0.756; **Table S1**) indicates a high level of overdispersion and significant deviation from a Poisson-distributed model. FACS data at high bulk MOI (open circles) were excluded from model fits due to the lack of confidence in high MOI measurements.

**Figure S3. MDCK cell survival patterns cannot be reproduced under a time-independent, input-dependent cell death rate model. (A)** The number of cells remaining for 3, 6, 12, and 18 hpi, respectively, as a function of bulk MOI, along with time-independent, input-dependent cell death rate model fits (lines). **(B)** Number of surviving MDCK cells that are infected at 18 hpi, as measured by FACS, along with the negative binomial distribution model fit (line). As in Figure 2C, statistical parameterization of this model (overdispersion parameter *r* = 0.756; **Table S1**) indicates a high level of overdispersion and significant deviation from a Poisson-distributed model. FACS data at high bulk MOI (open circles) were excluded from model fits due to the lack of confidence in high MOI measurements.

**Figure S4. Comparison of Poisson, zero-inflated Poisson, and negative binomial distribution fits to MDCK and A549 FACS data. (A)** Number of surviving MDCK cells infected at 18 hpi (dots) and viral dispersion model fits to these data (lines). Under the most supported cell death rate model (the time-dependent, input-independent model), the best fit to the FACS data occurred under the negative binomial model with an overdispersion parameter of *r* = 0.597 (solid orange line; **Table S1**). FACS data points from the high MOI experiments (open circles) were excluded from the model fit. Higher levels of overdispersion (*r* = 0.2; blue line) underestimated percentages of infected cells at 18 hpi. Lower levels of overdispersion (*r* = 2; blue line) overestimated percentages of infected cells at 18 hpi. To obtain the negative binomial models at fixed dispersion parameter values, *r* = 0.2, 2, we re-fit the parameters of the time-dependent, input-independent cell death rate model. A Poisson distribution assumption (r = ∞; solid red line) severely overestimated percentages of infected cells at 18 hpi. The zero-inflated Poisson is shown with the time-dependent, input-independent cell death rate model and with the probability of extra zeros, *p* = 0.312 (dashed red line). **Table S1** shows the four cell death rate models parameterized under the assumption of Poisson, negative binomial, and zero-inflated Poisson distributions for viral input across cells. ΔAIC values for these models are significantly larger than 0, indicating that the negative binomial distribution model is strongly preferred over both the Poisson and zero-inflated Poisson distribution models. **(B)** Number of surviving A549 cells infected at 18 hpi (dots) and viral dispersion model fits to these data (lines). Under the most supported cell death rate model (the time-dependent, input-independent model), the best fit to the FACS data occurred under the negative binomial model with an overdispersion parameter of *r* = 0.338 (solid orange line; **Table S2**). FACS data points from the high MOI experiments (open circles) were excluded from the model fit. Higher levels of overdispersion (*r* = 0.1; dashed blue line) underestimated percentages of infected cells at 18 hpi. Lower levels of overdispersion (*r* = 1; dashed blue line) overestimated percentages of infected cells at 18 hpi. A Poisson distribution assumption (r = ∞; solid red line) severely overestimated percentages of infected cells at 18 hpi. The zero-inflated Poisson is shown with the time-dependent, input-independent cell death rate model and with the probability of extra zeros, *p* = 0.493 (dashed red line). **Table S2** shows the four cell death rate models parameterized under the assumption of Poisson, negative binomial, and zero-inflated Poisson distributions for viral input across A549 cells. ΔAIC values for these models are significantly larger than 0, indicating that the negative binomial distribution model is also strongly preferred in A549 cells over the Poisson distribution models.

**Figure S5. Most supported time-independent models of virus production cannot capture virus production kinetics. (A)** Time-independent, linear input-dependent model fits to virus production in MDCK cells overestimate viral output at 6 hpi and underestimate the output at 18 hpi. **(B)** The virus production rate is constant over time and the rate increases linearly with increasing cellular MOI: i = 1, 4, 8. **(C)** Time-independent, input-independent model fits to virus production in A549 cells overestimate viral output at 6 hpi and underestimate viral output at 18 hpi. **(D)** The virus production rate is constant over time and independent of the cellular MOI.

**Figure S6. The input-independent model overestimates virus output at low bulk MOI and underestimates virus output at high bulk MOI in MDCK cells. (A)** The time delay in virus production was estimated in this model to be 5.27 days. After that point, the virus production rate was assumed to increase linearly in time, with an estimated slope of 2.52. **(B)** Model fits to virus production in MDCK cells (**Table S3**).

**Figure S7. The linear input-dependent model cannot capture virus production in A549 cells. (A)** The virus production rate is zero until ~ 5 hpi after which point the rate increases linearly in time. The slope of this linear increase depends on the cellular MOI: i=1, 4, 8 (**Table S4**). **(B)** The linear input-dependent model fits to data: at 6 and 12 hpi, the model overestimates viral output for high bulk MOI values, and at 18 hpi, the model underestimates viral output for low bulk MOI values.

**Figure S8. Cellular co-infection enhances ISG induction in A549 but not MDCK.** MDCK and A549 cells were infected with PR8 under single cycle conditions at the range of bulk MOIs: 0.08-7.86 and 0.06-26.1, respectively. Levels of cellular IFNB1 and IFNL1 transcript were measured by RT-qPCR at 8 and 18 hpi, compared to levels in mock cells. Interferon stimulated genes (ISGs) induction relative to mock vs. bulk MOI in MDCK cells at 8 and 18 hpi; ISG15, ZC3HAV1, and Mx1 did not show significant positive correlation between ISG induction and bulk MOI at 8 hpi (p = 0.69, p = 0.11, p = 0.46, respectively) nor 18 hpi (p = 0.06, p = 0.17, p = 0.08, respectively). ISGs induction relative to mock vs. bulk MOI in A549 cells at 8 and 18 hpi showed different temporal patterns of induction. Significant positive correlation between ISG15 induction and bulk MOI was not found at 8 hpi (p = 0.07) yet at 18 hpi (p = 0.0002). Significant positive correlation between ZC3HAV1 induction and bulk MOI was found at 8 hpi (p = 0.01) yet not at 18 hpi (p = 0.31). For Mx1, there is a significant positive correlation to bulk MOI at both timepoints (p = 0.02 for both).

## SUPPLEMENTAL TABLE LEGENDS

**Table S1. Fits of cell death rate models to MDCK cell data.** Rows correspond to distinct cell death rate models. The mathematical formulation for each cell death rate model is provided in the second column. The models are group by the virus distribution assumption, going from top to bottom: Poisson, Negative binomial, zero-inflated Poisson. Point estimates and 95% confidence intervals are provided in the third column for each model’s parameters. Confidence intervals for parameter estimates close to zero were omitted (Methods). Units of the parameters are provided in the fourth column. The fifth column lists the residual sum of squares (RSS) for each model, parameterized with the point estimates of the third column. The model most supported by the data is the time-dependent, input-independent model (ΔAIC = 0). Models with higher ΔAIC have less statistical support.

**Table S2. Fits of cell death rate models to A549 cell data.** As in Table S1, rows correspond to distinct cell death rate models assuming the viral infection distribution from top to bottom: Poisson, negative binomial, zero-inflated Poisson. The model most supported by the data is the time-dependent, input-independent model (ΔAIC = 0).

**Table S3. Fits of viral production rate models to MDCK cell data.** Rows correspond to distinct viral production rate models. Parameter estimates are given along with 95 percent confidence intervals for viral production rate model parameters. The model that is most supported by the data has ΔAIC = 0, and models with higher ΔAIC have less statistical support. Confidence intervals for high parameter estimates were omitted (see methods).

**Table S4. Fits of viral production rate models to A549 cell data.** Rows correspond to distinct viral production rate models. Parameter estimates are given along with 95 percent confidence intervals for viral production rate model parameters. The model that is most supported by the data has ΔAIC = 0, and models with higher ΔAIC have less statistical support.

**Table S5. Descriptions of interferon induction models.** We considered three IFN induction models in which the induction rates are independent of time but differ based on the viral input. We fit an input-independent, linear input-dependent, and saturating input-dependent induction rate models to data at 8 and 18 hpi, giving a total of nine model combinations (see Table S6).

**Table S6. Fits of IFN induction models to IFNL1 data in A549 cells.** Rows correspond to different combinations of the IFN induction rate models listed in Table 5 from 0-8 hpi and from 8-18 hpi. The first column gives the model number from 0-8 hpi, and the third column gives the model number from 8-18 hpi. Parameter estimates are given along with 95 percent confidence intervals for IFN induction rate model parameters. The model that is most supported by the data has ΔAIC = 0, and models with higher ΔAIC have less statistical support. Confidence intervals for high parameter estimates were omitted (see methods).

**Table S7. Fits of superinfection exclusion models to FACs data in MDCK cells.** Rows correspond to distinct viral production rate models. The first model assumed that all rH3N1-infected cells had the same reduced probability of becoming infected with rH1N2 (input-independent). The second model assumed that the probability of being infected with rH1N2 decreased with cellular rH3N1 MOI (input-dependent). Parameter estimates are given along with 95 percent confidence intervals for viral production rate model parameters. The model that is most supported by the data has ΔAIC = 0, and models with higher ΔAIC have less statistical support.

**Table S8. RT-qPCR primers for the quantification of interferons, interferon stimulated genes, and endogenous control for MDCK (canine) and A549 (human) cell lines.** SYBR green primers were used for the quantification of canine targets of IFNB1, IFNL1, ISG15, and Mx1, with β-actin as the endogenous control. For A549, Taqman assays were used for the quantification of IFNB1 and IFNL1 with GADPH as the endogenous control and SYBR green chemistry was used for the quantification of ISG15 and Mx1, with β-actin as the endogenous control.

## SUPPLEMENTAL SOURCE DATA FILE LEGEND

This file contains the source data used to generate every figure (main and supplemental) in this manuscript. Each tab of the Excel file includes figure panels, which are often grouped according to cell line, MDCK or A549 cells. For data that was used in multiple figures, we included these only once and made a note within the sheet of any other figures that also show these data. Oftentimes, the data are included in both a main figure (found in the text) and one or more supplemental figures; in these cases, we labeled the tabs according to the main figure.

## Notes

### Competing Interest Statement

The authors have declared no competing interest.

### Summary of Updates

Added new experimental data and analyses

