## Supplemental figures for "Cellular co-infection can modulate the efficiency of influenza A virus production and shape the interferon response"

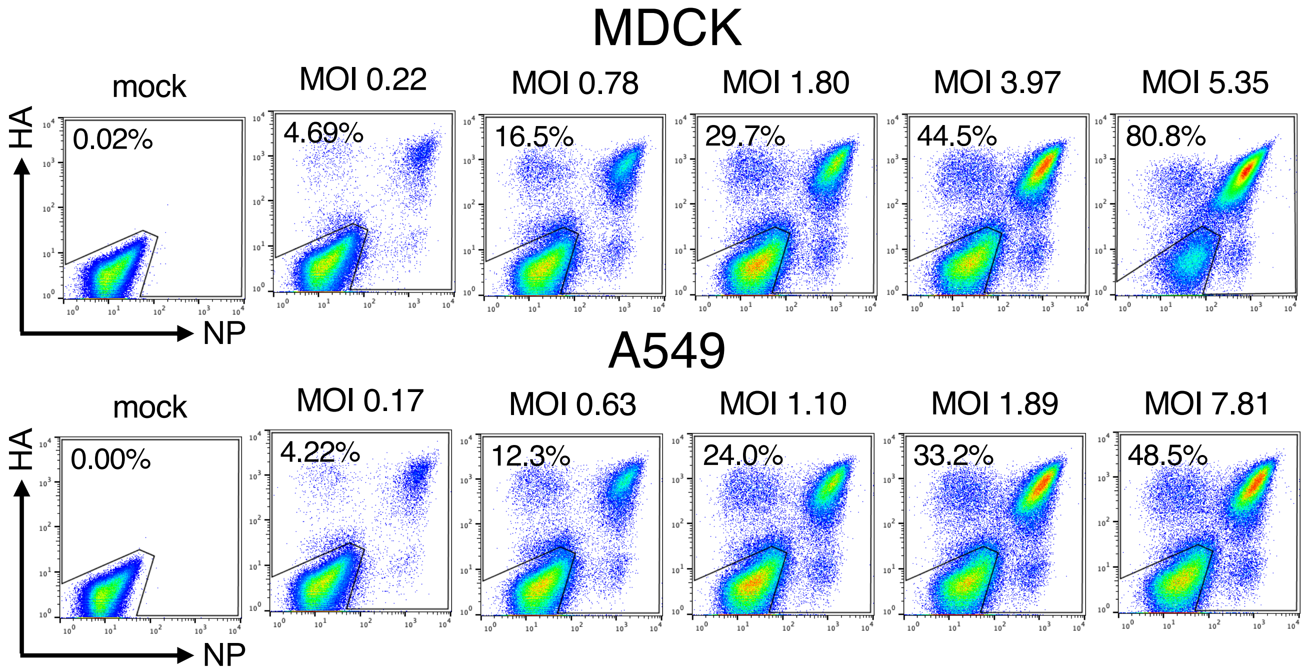


**Figure S1. Fluorescence-activated cell sorting based on HA and NP expression** for MDCK (top row) and A549 (bottom row) cells. Mock infected cells were used for the gating of all cells that are productively infected and positive for the expression of either HA or NP. For MDCK at the intended MOI of 10. A modified gating was used for the MDCK cell line at the MOI of 5.35 for MDCK and 7.81 for A549 to more effectively exclude negative cells.

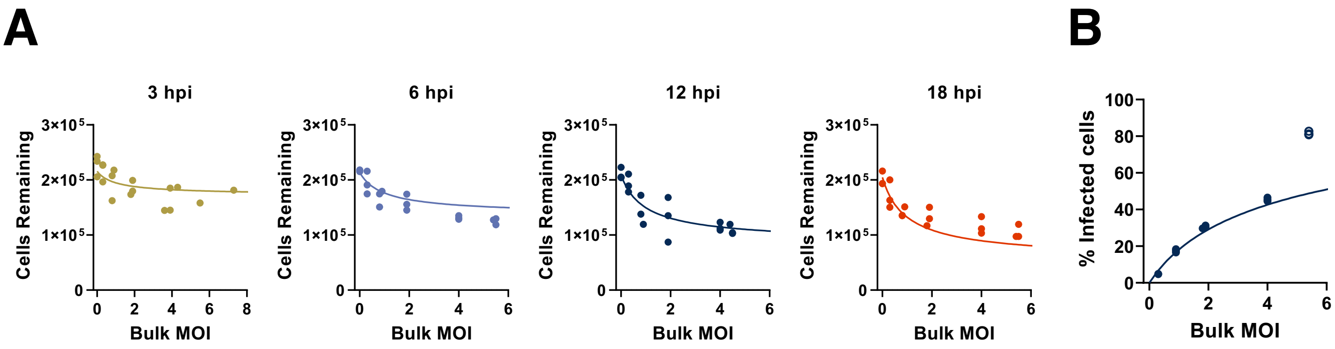


**Figure S2. MDCK cell survival patterns cannot be reproduced under a time-independent, input-independent cell death rate model. (A)** The number of cells remaining for 3, 6, 12, and 18 hpi, respectively, as a function of bulk MOI, along with time-independent, input-independent cell death rate model fits (lines). **(B)** Number of surviving MDCK cells that are infected at 18 hpi, as measured by FACS, along with the negative binomial distribution model fit (line). As in Figure 2C, statistical parameterization of this model (overdispersion parameter $r = 0.756$; **Table S1**) indicates a high level of overdispersion and significant deviation from a Poisson-distributed model. FACS data at high bulk MOI (open circles) were excluded from model fits due to the lack of confidence in high MOI measurements.

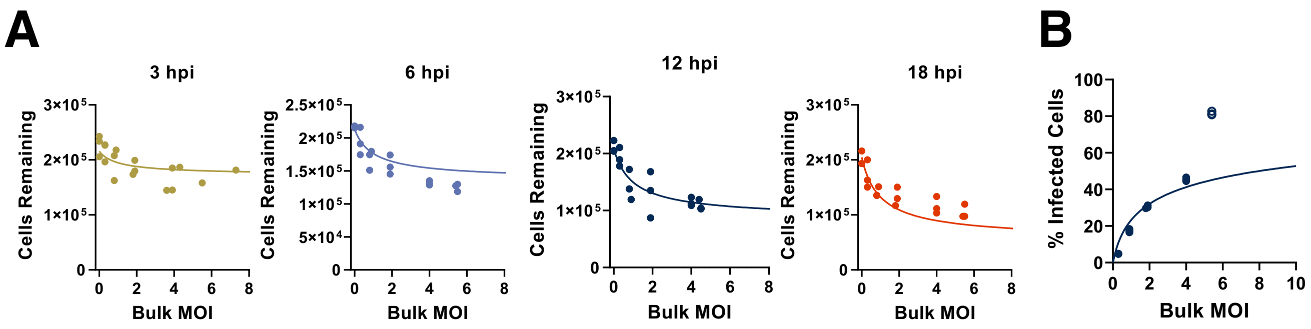


**Figure S3. MDCK cell survival patterns cannot be reproduced under a time-independent, input-dependent cell death rate model. (A)** The number of cells remaining for 3, 6, 12, and 18 hpi, respectively, as a function of bulk MOI, along with time-independent, input-dependent cell death rate model fits (lines). **(B)** Number of surviving MDCK cells that are infected at 18 hpi, as measured by FACS, along with the negative binomial distribution model fit (line). As in Figure 2C, statistical parameterization of this model (overdispersion parameter $r = 0.756$; **Table S1**) indicates a high level of overdispersion and significant deviation from a Poisson-distributed model. FACS data at high bulk MOI (open circles) were excluded from model fits due to the lack of confidence in high MOI measurements.

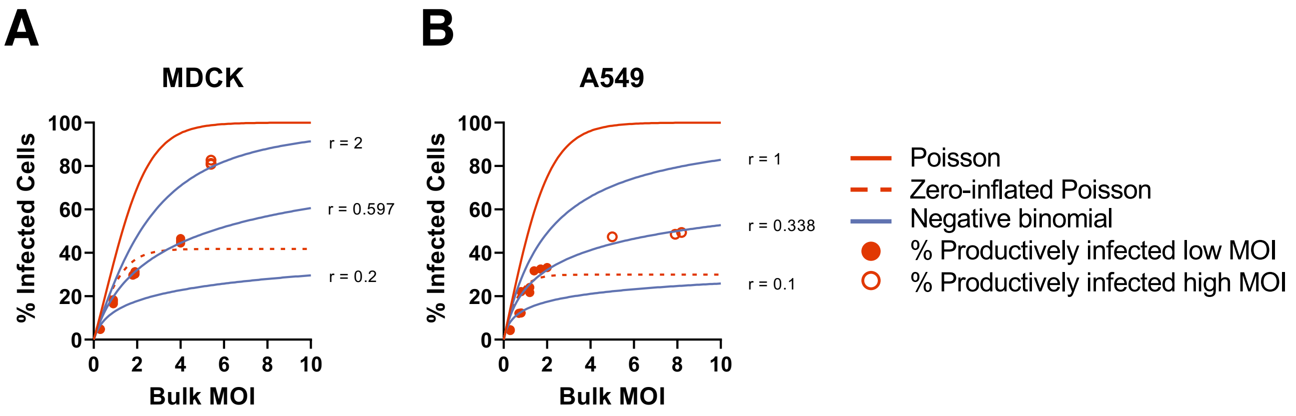


**Figure S4. Comparison of Poisson, zero-inflated Poisson, and negative binomial distribution fits to MDCK and A549 FACS data. (A)** Number of surviving MDCK cells infected at 18 hpi (dots) and viral dispersion model fits to these data (lines). Under the most supported cell death rate model (the time-dependent, input-independent model), the best fit to the FACS data occurred under the negative binomial model with an overdispersion parameter of $r = 0.597$ (solid orange line; **Table S1**). FACS data points from the high MOI experiments (open circles) were excluded from the model fit. Higher levels of overdispersion ($r$ = 0.2; blue line) underestimated percentages of infected cells at 18 hpi. Lower levels of overdispersion ($r$ = 2; blue line) overestimated percentages of infected cells at 18 hpi. To obtain the negative binomial models at fixed dispersion parameter values, $r$ = 0.2, 2, we re-fit the parameters of the time-dependent, input-independent cell death rate model. A Poisson distribution assumption (r = ∞; solid red line) severely overestimated percentages of infected cells at 18 hpi. The zero-inflated Poisson is shown with the time-dependent, input-independent cell death rate model and with the probability of extra zeros, $p$ = 0.312 (dashed red line). **Table S1** shows the four cell death rate models parameterized under the assumption of Poisson, negative binomial, and zero-inflated Poisson distributions for viral input across cells. ΔAIC values for these models are significantly larger than 0, indicating that the negative binomial distribution model is strongly preferred over both the Poisson and zero-inflated Poisson distribution models. **(B)** Number of surviving A549 cells infected at 18 hpi (dots) and viral dispersion model fits to these data (lines). Under the most supported cell death rate model (the time-dependent, input-independent model), the best fit to the FACS data occurred under the negative binomial model with an overdispersion parameter of $r = 0.338$ (solid orange line; **Table S2**). FACS data points from the high MOI experiments (open circles) were excluded from the model fit. Higher levels of overdispersion ($r = 0.1$; dashed blue line) underestimated percentages of infected cells at 18 hpi. Lower levels of overdispersion ($r$ = 1; dashed blue line) overestimated percentages of infected cells at 18 hpi. A Poisson distribution assumption (r = ∞; solid red line) severely overestimated percentages of infected cells at 18 hpi. The zero-inflated Poisson is shown with the time-dependent, input-independent cell death rate model and with the probability of extra zeros, $p$ = 0.493 (dashed red line). **Table S2** shows the four cell death rate models parameterized under the assumption of Poisson, negative binomial, and zero-inflated Poisson distributions for viral input across A549 cells. ΔAIC values for these models are significantly larger than 0, indicating that the negative binomial distribution model is also strongly preferred in A549 cells over the Poisson distribution models.

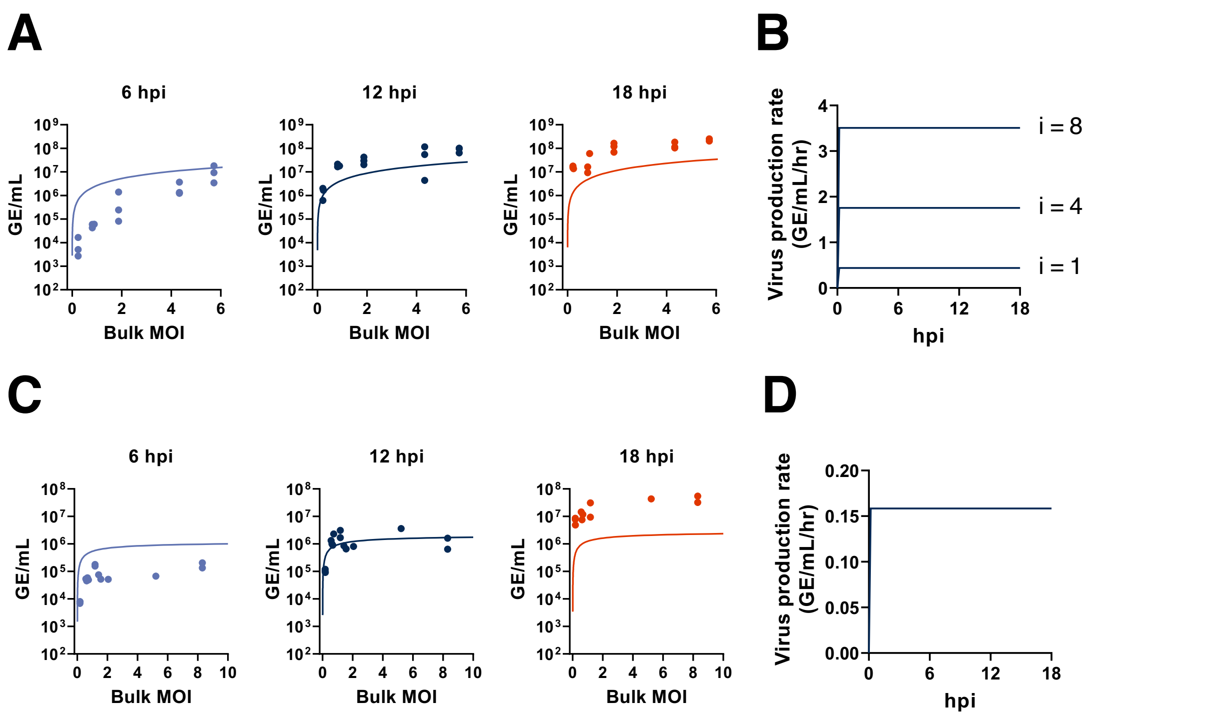


**Figure S5. Most supported time-independent models of virus production cannot capture virus production kinetics. (A)** Time-independent, linear input-dependent model fits to virus production in MDCK cells overestimate viral output at 6 hpi and underestimate the output at 18 hpi. **(B)** The virus production rate is constant over time and the rate increases linearly with increasing cellular MOI: i = 1, 4, 8. **(C)** Time-independent, input-independent model fits to virus production in A549 cells overestimate viral output at 6 hpi and underestimate viral output at 18 hpi. **(D)** The virus production rate is constant over time and independent of the cellular MOI.

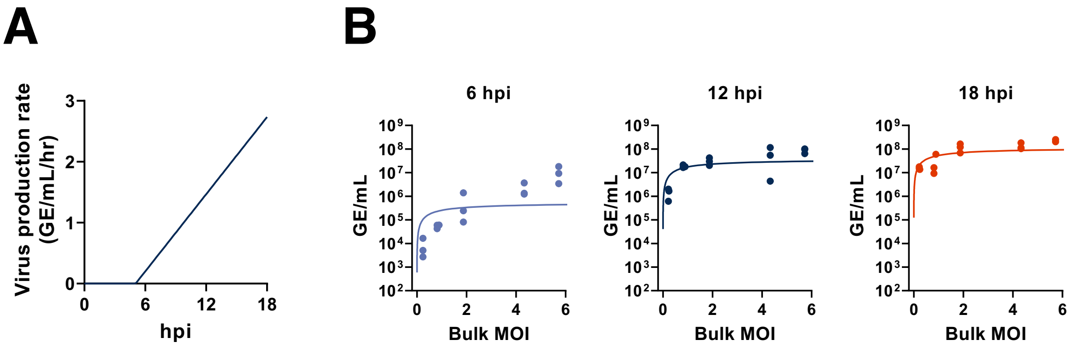


**Figure S6. The input-independent model overestimates virus output at low bulk MOI and underestimates virus output at high bulk MOI in MDCK cells. (A)** The time delay in virus production was estimated in this model to be 5.27 days. After that point, the virus production rate was assumed to increase linearly in time, with an estimated slope of 2.52. **(B)** Model fits to virus production in MDCK cells (**Table S3**).

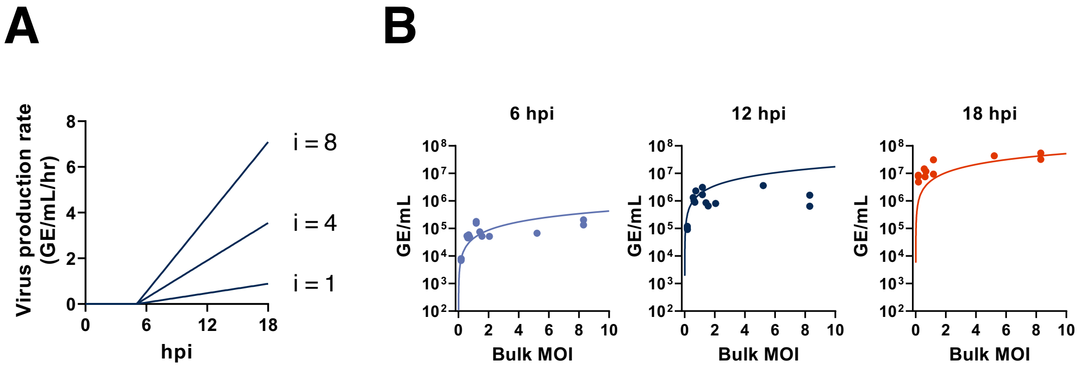


**Figure S7. The linear input-dependent model cannot capture virus production in A549 cells. (A)** The virus production rate is zero until ~ 5 hpi after which point the rate increases linearly in time. The slope of this linear increase depends on the cellular MOI: i=1, 4, 8. **(B)** The linear input-dependent model fits to data: at 6 and 12 hpi, the model overestimates viral output for high bulk MOI values, and at 18 hpi, the model underestimates viral output for low bulk MOI values.

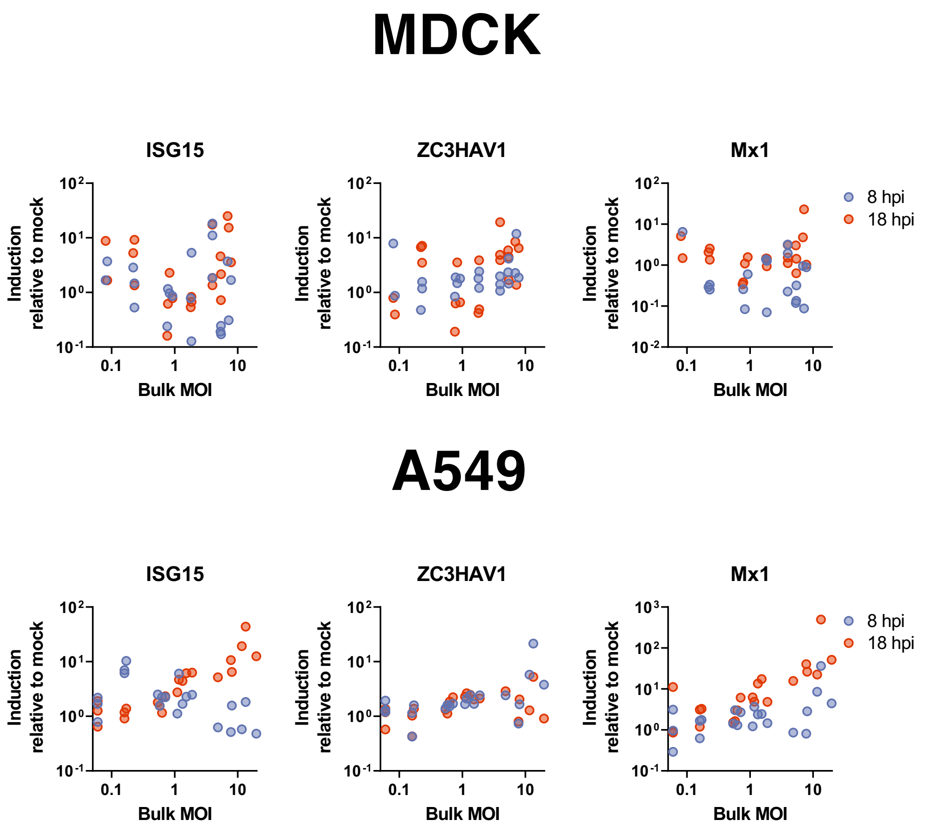


**Figure S8. Cellular co-infection enhances ISG induction in A549 but not MDCK.** MDCK and A549 cells were infected with PR8 under single cycle conditions at the range of bulk MOIs: 0.08-7.86 and 0.06-26.1, respectively. Levels of cellular IFNB1 and IFNL1 transcript were measured by RT-qPCR at 8 and 18 hpi, compared to levels in mock cells. Interferon stimulated genes (ISGs) induction relative to mock vs. bulk MOI in MDCK cells at 8 and 18 hpi; ISG15, ZC3HAV1, and Mx1 did not show significant positive correlation between ISG induction and bulk MOI at 8 hpi ($p = 0.69$, $p = 0.11$, $p = 0.46$, respectively) nor 18 hpi ($p = 0.06$, $p = 0.17$, $p = 0.08$, respectively). ISGs induction relative to mock vs. bulk MOI in A549 cells at 8 and 18 hpi showed different temporal patterns of induction. Significant positive correlation between ISG15 induction and bulk MOI was not found at 8 hpi ($p = 0.07$) yet at 18 hpi ($p = 0.0002$). Significant positive correlation between ZC3HAV1 induction and bulk MOI was found at 8 hpi ($p = 0.01$) yet not at 18 hpi ($p = 0.31$). For Mx1, there is a significant positive correlation to bulk MOI at both timepoints ($p = 0.02$ for both).
