## Supplemental tables for "Cellular co-infection can modulate the efficiency of influenza A virus production and shape the interferon response"

#### Table S1. Fits of cell death rate models to MDCK cell data.

| Cell death rate model | Cell death rate $\left(\mathrm{hr}^{-1}\right)$ | Parameter estimates (95% CI) | Units | RSS | ΔAIC | Parameter legend |
|---|---|---|---|---|---|---|
| Time-independent,<br>input-independent death rate | $\alpha(s,i) = a$ | $a = 0.0665$ $[0.0537, 0.0823]$<br>$\mu = 0.0025$ $[0.0003, 0.0189]$<br>$N_{cells} = 225,000$ $[214,000, 237,000]$ | $\mathrm{hr}^{-1}$<br>$\mathrm{hr}^{-1}$ | 18,600 | 133.3 | $a$ = infected cell death rate<br>$\mu$ = constant background cell death rate<br>$N_{cells}$ = number of cells |
| Time-dependent,<br>input-independent death rate | $\alpha(s,i) = b\,k\,s^{k-1}$ | $b = 0.208$ $[0.127, 0.339]$<br>$k = 0.546$ $[0.344, 0.867]$<br>$\mu = 0.0092$ $[0.0043, 0.0195]$<br>$N_{cells} = 247,000$ $[230,000, 265,000]$ | $\mathrm{hr}^{-k}$<br><br>$\mathrm{hr}^{-1}$ | 17,100 | 128.1 | $b$ = scale parameter<br>$k$ = shape parameter<br>$\mu$ = constant background cell death rate<br>$N_{cells}$ = number of cells |
| Time-independent,<br>input-dependent death rate | $\alpha(s,i) = c\,i^{\varepsilon}$ | $c = 0.0665$ $[0.0537, 0.0823]$<br>$\varepsilon = 2.62 \times 10^{-15}$ $[-,-]$<br>$\mu = 0.0025$ $[0.0003, 0.0189]$<br>$N_{cells} = 225,000$ $[214,000, 237,000]$ | $\mathrm{hr}^{-1}\cdot$ virion$^{-\varepsilon}$<br><br>$\mathrm{hr}^{-1}$ | 18,600 | 135.3 | $c$ = effective per virion infected cell death rate<br>$\varepsilon$ = exponent<br>$\mu$ = constant background cell death rate<br>$N_{cells}$ = number of cells |
| Time-dependent,<br>input-dependent death rate | $\alpha(s,i) = d\,k\,s^{k-1}\,i^{\varepsilon}$ | $d = 0.208$ $[0.127, 0.339]$<br>$k = 0.546$ $[0.345, 0.867]$<br>$\varepsilon = 1.67 \times 10^{-15}$ $[-,-]$<br>$\mu = 0.0092$ $[0.0043, 0.0195]$<br>$N_{cells} = 247,000$ $[230,000, 265,000]$ | $\mathrm{hr}^{-k}\cdot$ virion$^{-\varepsilon}$<br><br><br>$\mathrm{hr}^{-1}$ | 17,100 | 130.1 | $d$ = effective per virion scale parameter<br>$k$ = shape parameter<br>$\varepsilon$ = exponent<br>$\mu$ = constant background cell death rate<br>$N_{cells}$ = number of cells |
| Time-independent,<br>input-independent death rate | $\alpha(s,i) = a$ | $a = 0.0767$ $[0.0692, 0.0851]$<br>$r = 0.756$ $[0.623, 0.917]$<br>$\mu = 0.0035$ $[0.0010, 0.0126]$<br>$N_{cells} = 218,000$ $[208,000, 228,000]$ | $\mathrm{hr}^{-1}$<br><br>$\mathrm{hr}^{-1}$ | 6,050 | 40.9 | $a$ = infected cell death rate<br>$r$ = dispersion parameter<br>$\mu$ = constant background cell death rate<br>$N_{cells}$ = number of cells |
| Time-dependent,<br>input-independent death rate | $\alpha(s,i) = b\,k\,s^{k-1}$ | $b = 0.447$ $[0.267, 0.748]$<br>$k = 0.308$ $[0.158, 0.600]$<br>$r = 0.597$ $[0.508, 0.703]$<br>$\mu = 0.0146$ $[0.0099, 0.0215]$<br>$N_{cells} = 250,000$ $[235,000, 266,000]$ | $\mathrm{hr}^{-k}$<br><br><br>$\mathrm{hr}^{-1}$ | 3,630 | 0 | $b$ = scale parameter<br>$k$ = shape parameter<br>$r$ = dispersion parameter<br>$\mu$ = constant background cell death rate<br>$N_{cells}$ = number of cells |
| Time-independent,<br>input-dependent death rate | $\alpha(s,i) = c\,i^{\varepsilon}$ | $c = 0.0767$ $[0.0692, 0.0851]$<br>$\varepsilon = 1.23 \times 10^{-14}$ $[-,-]$<br>$r = 0.756$ $[0.623, 0.917]$<br>$\mu = 0.0035$ $[0.0010, 0.0126]$<br>$N_{cells} = 218,000$ $[208,000, 228,000]$ | $\mathrm{hr}^{-1}\cdot$ virion$^{-\varepsilon}$<br><br><br>$\mathrm{hr}^{-1}$ | 6,050 | 42.9 | $c$ = effective per virion infected cell death rate<br>$\varepsilon$ = exponent<br>$r$ = dispersion parameter<br>$\mu$ = constant background cell death rate<br>$N_{cells}$ = number of cells |
| Time-dependent,<br>input-dependent death rate | $\alpha(s,i) = d\,k\,s^{k-1}\,i^{\varepsilon}$ | $d = 0.447$ $[0.267, 0.748]$<br>$k = 0.308$ $[0.158, 0.600]$<br>$\varepsilon = 2.70 \times 10^{-15}$ $[-,-]$<br>$r = 0.597$ $[0.508, 0.703]$<br>$\mu = 0.0146$ $[0.0099, 0.0215]$<br>$N_{cells} = 250,000$ $[235,000, 266,000]$ | $\mathrm{hr}^{-k}\cdot$ virion$^{-\varepsilon}$<br><br><br><br>$\mathrm{hr}^{-1}$ | 3,630 | 2.0 | $d$ = effective per virion scale parameter<br>$k$ = shape parameter<br>$\varepsilon$ = exponent<br>$r$ = dispersion parameter<br>$\mu$ = constant background cell death rate<br>$N_{cells}$ = number of cells |
| Time-independent,<br>input-independent death rate | $\alpha(s,i) = a$ | $a = 0.0805$ $[0.0725, 0.0894]$<br>$p = 0.264$ $[0.223, 0.312]$<br>$\mu = 0.0025$ $[0.0004, 0.0156]$<br>$N_{cells} = 220,000$ $[210,000, 230,000]$ | $\mathrm{hr}^{-1}$<br><br>$\mathrm{hr}^{-1}$ | 6,270 | 43.9 | $a$ = infected cell death rate<br>$p$ = probability extra zeros<br>$\mu$ = constant background cell death rate<br>$N_{cells}$ = number of cells |
| Time-dependent,<br>input-independent death rate | $\alpha(s,i) = b\,k\,s^{k-1}$ | $b = 0.439$ $[0.259, 0.743]$<br>$k = 0.342$ $[0.186, 0.628]$<br>$p = 0.312$ $[0.271, 0.360]$<br>$\mu = 0.0137$ $[0.0089, 0.0210]$<br>$N_{cells} = 253,000$ $[236,000, 272,000]$ | $\mathrm{hr}^{-k}$<br><br><br>$\mathrm{hr}^{-1}$ | 4,010 | 8.4 | $b$ = scale parameter<br>$k$ = shape parameter<br>$p$ = probability extra zeros<br>$\mu$ = constant background cell death rate<br>$N_{cells}$ = number of cells |
| Time-independent,<br>input-dependent death rate | $\alpha(s,i) = c\,i^{\varepsilon}$ | $c = 0.0704$ $[0.0538, 0.0921$<br>$\varepsilon = 0.00103$ $[0.0161, 0.658]$<br>$p = 0.272$ $[0.224, 0.332]$<br>$\mu = 0.0028$ $[0.0006, 0.0139]$<br>$N_{cells} = 219,000$ $[209,000, 229,000]$ | $\mathrm{hr}^{-1}\cdot$ virion$^{-\varepsilon}$<br><br><br>$\mathrm{hr}^{-1}$ | 6,200 | 45.0 | $c$ = effective per virion infected cell death rate<br>$\varepsilon$ = exponent<br>$p$ = probability extra zeros<br>$\mu$ = constant background cell death rate<br>$N_{cells}$ = number of cells |
| Time-dependent,<br>input-dependent death rate | $\alpha(s,i) = d\,k\,s^{k-1}\,i^{\varepsilon}$ | $d = 0.418$ $[0.235, 0.745]$<br>$k = 0.345$ $[0.189, 0.629]$<br>$\varepsilon = 0.0339$ $[0.000380, 3.03]$<br>$p = 0.3146$ $[0.0088, 0.0209]$<br>$\mu = 0.0136$ $[0.0088, 0.0209]$<br>$N_{cells} = 252,000$ $[235,000, 271,000]$ | $\mathrm{hr}^{-k}\cdot$ virion$^{-\varepsilon}$<br><br><br><br>$\mathrm{hr}^{-1}$ | 4,000 | 10.3 | $d$ = effective per virion scale parameter<br>$k$ = shape parameter<br>$\varepsilon$ = exponent<br>$p$ = probability extra zeros<br>$\mu$ = constant background cell death rate<br>$N_{cells}$ = number of cells |

**Table S1. Fits of cell death rate models to MDCK cell data.** Rows correspond to distinct cell death rate models. The mathematical formulation for each cell death rate model is provided in the second column. The models are group by the virus distribution assumption, going from top to bottom: Poisson, Negative binomial, zero-inflated Poisson. Point estimates and 95% confidence intervals are provided in the third column for each model's parameters. Confidence intervals for parameter estimates close to zero were omitted

(Methods). Units of the parameters are provided in the fourth column. The fifth column lists the residual sum of squares (RSS) for each model, parameterized with the point estimates of the third column. The model most supported by the data is the time-dependent, input-independent model ($\Delta$AIC = 0). Models with higher $\Delta$AIC have less statistical support.

### Table S2. Fits of cell death rate models to A549 cell data.

| Cell death rate model | Cell death rate $\left(\mathrm{hr}^{-1}\right)$ | Parameter estimates (95% CI) | | Units | RSS | ΔAIC | Parameter legend |
|---|---|---|---|---|---|---|---|
| Time-independent,<br>input-independent death rate | $\alpha(s,i) = a$ | $a = 0.0405$<br>$\mu = 0.0052$<br>$N_{cells} = 250,000$ | $[0.0297, 0.0552]$<br>$[0.0024, 0.0112]$<br>$[242,000, 258,000]$ | $\mathrm{hr}^{-1}$<br>$\mathrm{hr}^{-1}$ | 9,840 | 110.2 | $a$ = infected cell death rate<br>$\mu$ = constant background cell death rate<br>$N_{cells}$ = number of cells |
| Time-dependent,<br>input-independent death rate | $\alpha(s,i) = b\,k\,s^{k-1}$ | $b = 0.0218$<br>$k = 1.23$<br>$\mu = 0.0036$<br>$N_{cells} = 243,000$ | $[0.0062, 0.0771]$<br>$[0.797, 1.89]$<br>$[0.0007, 0.0187]$<br>$[232,000, 255,000]$ | $\mathrm{hr}^{-k}$<br><br>$\mathrm{hr}^{-1}$ | 9,760 | 111.6 | $b$ = scale parameter<br>$k$ = shape parameter<br>$\mu$ = constant background cell death rate<br>$N_{cells}$ = number of cells |
| Time-independent,<br>input-dependent death rate | $\alpha(s,i) = c\,i^{\varepsilon}$ | $c = 0.0405$<br>$\varepsilon = 3.65 \times 10^{-16}$<br>$\mu = 0.0052$<br>$N_{cells} = 250,000$ | $[0.0297, 0.0552]$<br>$[-,-]$<br>$[0.0024, 0.0112]$<br>$[242,000, 258,000]$ | $\mathrm{hr}^{-1} \cdot \mathrm{virion}^{-\varepsilon}$<br><br>$\mathrm{hr}^{-1}$ | 9840 | 112.2 | $c$ = effective per virion infected cell death rate<br>$\varepsilon$ = exponent<br>$\mu$ = constant background cell death rate<br>$N_{cells}$ = number of cells |
| Time-dependent,<br>input-dependent death rate | $\alpha(s,i) = d\,k\,s^{k-1}\,i^{\varepsilon}$ | $d = 0.0218$<br>$k = 1.23$<br>$\varepsilon = 2.12 \times 10^{-15}$<br>$\mu = 0.0036$<br>$N_{cells} = 243,000$ | $[0.0062, 0.0771]$<br>$[0.798, 1.89]$<br>$[-,-]$<br>$[0.0007, 0.0187]$<br>$[232,000, 255,000]$ | $\mathrm{hr}^{-k} \cdot \mathrm{virion}^{-\varepsilon}$<br><br><br>$\mathrm{hr}^{-1}$ | 9,760 | 113.6 | $d$ = effective per virion scale parameter<br>$k$ = shape parameter<br>$\varepsilon$ = exponent<br>$\mu$ = constant background cell death rate<br>$N_{cells}$ = number of cells |
| Time-independent,<br>input-independent death rate | $\alpha(s,i) = a$ | $a = 0.0412$<br>$r = 0.3758$<br>$\mu = 0.0088$<br>$N_{cells} = 244,000$ | $[0.0349, 0.0485]$<br>$[0.3000, 0.4707]$<br>$[0.0063, 0.0123]$<br>$[239,000, 250,000]$ | $\mathrm{hr}^{-1}$<br><br>$\mathrm{hr}^{-1}$ | 2,660 | 2.3 | $a$ = infected cell death rate<br>$r$ = dispersion parameter<br>$\mu$ = constant background cell death rate<br>$N_{cells}$ = number of cells |
| Time-dependent,<br>input-independent death rate | $\alpha(s,i) = b\,k\,s^{k-1}$ | $b = 0.118$<br>$k = 0.601$<br>$r = 0.338$<br>$\mu = 0.0118$<br>$N_{cells} = 254,000$ | $[0.0529, 0.2632]$<br>$[0.362, 0.996]$<br>$[0.270, 0.425]$<br>$[0.0083, 0.0169]$<br>$[242,000, 265,000]$ | $\mathrm{hr}^{-k}$<br><br><br>$\mathrm{hr}^{-1}$ | 2,530 | 0 | $b$ = scale parameter<br>$k$ = shape parameter<br>$r$ = dispersion parameter<br>$\mu$ = constant background cell death rate<br>$N_{cells}$ = number of cells |
| Time-independent,<br>input-dependent death rate | $\alpha(s,i) = c\,i^{\varepsilon}$ | $c = 0.0412$<br>$\varepsilon = 1.35 \times 10^{-14}$<br>$r = 0.3758$<br>$\mu = 0.0088$<br>$N_{cells} = 244,000$ | $[0.0350, 0.0485]$<br>$[-,-]$<br>$[0.3000, 0.4708]$<br>$[0.0063, 0.0123]$<br>$[239,000, 250,000]$ | $\mathrm{hr}^{-1} \cdot \mathrm{virion}^{-\varepsilon}$<br><br><br>$\mathrm{hr}^{-1}$ | 2,660 | 4.3 | $c$ = effective per virion infected cell death rate<br>$\varepsilon$ = exponent<br>$r$ = dispersion parameter<br>$\mu$ = constant background cell death rate<br>$N_{cells}$ = number of cells |
| Time-dependent,<br>input-dependent death rate | $\alpha(s,i) = d\,k\,s^{k-1}\,i^{\varepsilon}$ | $d = 0.118$<br>$k = 0.601$<br>$\varepsilon = 2.02 \times 10^{-12}$<br>$r = 0.338$<br>$\mu = 0.0118$<br>$N_{cells} = 254,000$ | $[0.0529, 0.263]$<br>$[0.362, 0.996]$<br>$[-,-]$<br>$[0.270, 0.425]$<br>$[0.0083, 0.0169]$<br>$[242,000, 265,000]$ | $\mathrm{hr}^{-k} \cdot \mathrm{virion}^{-\varepsilon}$<br><br><br><br>$\mathrm{hr}^{-1}$ | 2,530 | 2.0 | $d$ = effective per virion scale parameter<br>$k$ = shape parameter<br>$\varepsilon$ = exponent<br>$r$ = dispersion parameter<br>$\mu$ = constant background cell death rate<br>$N_{cells}$ = number of cells |
| Time-independent,<br>input-independent death rate | $\alpha(s,i) = a$ | $a = 0.0499$<br>$p = 0.4746$<br>$\mu = 0.0074$<br>$N_{cells} = 246,000$ | $[0.0425, 0.0586]$<br>$[0.424, 0.532]$<br>$[0.0049, 0.0111]$<br>$[240,000, 251,000]$ | $\mathrm{hr}^{-1}$<br><br>$\mathrm{hr}^{-1}$ | 2,760 | 5.6 | $a$ = infected cell death rate<br>$p$ = probability extra zeros<br>$\mu$ = constant background cell death rate<br>$N_{cells}$ = number of cells |
| Time-dependent,<br>input-independent death rate | $\alpha(s,i) = b\,k\,s^{k-1}$ | $b = 0.1079$<br>$k = 0.7111$<br>$p = 0.493$<br>$\mu = 0.0098$<br>$N_{cells} = 253,000$ | $[0.0387, 0.301]$<br>$[0.413, 1.22]$<br>$[0.436, 0.556]$<br>$[0.0061, 0.0159]$<br>$[240,000, 268,000]$ | $\mathrm{hr}^{-k}$<br><br><br>$\mathrm{hr}^{-1}$ | 2,690 | 5.4 | $b$ = scale parameter<br>$k$ = shape parameter<br>$p$ = probability extra zeros<br>$\mu$ = constant background cell death rate<br>$N_{cells}$ = number of cells |
| Time-independent,<br>input-dependent death rate | $\alpha(s,i) = c\,i^{\varepsilon}$ | $c = 0.0704$<br>$\varepsilon = 0.136$<br>$p = 0.5017$<br>$\mu = 0.0078$<br>$N_{cells} = 245,000$ | $[0.0538, 0.0921]$<br>$[0.0328, 0.564]$<br>$[0.432, 0.583]$<br>$[0.0054, 0.0114]$<br>$[240,000, 251,000]$ | $\mathrm{hr}^{-1} \cdot \mathrm{virion}^{-\varepsilon}$<br><br><br>$\mathrm{hr}^{-1}$ | 2,690 | 5.4 | $c$ = effective per virion infected cell death rate<br>$\varepsilon$ = exponent<br>$p$ = probability extra zeros<br>$\mu$ = constant background cell death rate<br>$N_{cells}$ = number of cells |
| Time-dependent,<br>input-dependent death rate | $\alpha(s,i) = d\,k\,s^{k-1}\,i^{\varepsilon}$ | $d = 0.108$<br>$k = 0.711$<br>$\varepsilon = 1.47 \times 10^{-9}$<br>$p = 0.493$<br>$\mu = 0.0098$<br>$N_{cells} = 253,000$ | $[0.0388, 0.301]$<br>$[0.413, 1.22]$<br>$[-,-]$<br>$[0.436, 0.556]$<br>$[0.0061, 0.0159]$<br>$[240,000, 268,000]$ | $\mathrm{hr}^{-k} \cdot \mathrm{virion}^{-\varepsilon}$<br><br><br><br>$\mathrm{hr}^{-1}$ | 2,690 | 7.4 | $d$ = effective per virion scale parameter<br>$k$ = shape parameter<br>$\varepsilon$ = exponent<br>$p$ = probability extra zeros<br>$\mu$ = constant background cell death rate<br>$N_{cells}$ = number of cells |

**Table S2. Fits of cell death rate models to A549 cell data.** As in Table 1, rows correspond to distinct cell death rate models assuming the viral infection distribution from top to bottom: Poisson, negative binomial, zero-inflated Poisson. The model most supported by the data is the time-dependent, input-independent model (ΔAIC = 0).

### Table S3. Fits of viral production rate models to MDCK cell data.

| **Time-independent production rate models** | **Viral production rate** $\left(\mathbf{hr}^{-1} \cdot \textbf{infected cell}^{-1}\right)$ | **Parameter estimates (95% CI)** | **Units** | **RSS** | **$\Delta$AIC** | **Parameter legend** |
|---|---|---|---|---|---|---|
| Input-independent | $v(t,i) = l$ | $l = 1.38$ [0.663, 2.88] | $\text{hr}^{-1} \cdot \text{infected cell}^{-1}$ | 272 | 69.2 | $l$ = viral production rate |
| Linear input-dependent | $v(t,i) = r\,i$ | $r = 0.438$ [0.223, 0.862] | $\text{hr}^{-1} \cdot \text{virion}^{-1} \cdot \text{infected cell}^{-1}$ | 231 | 62.0 | $r$ = per virion production rate |
| Saturating input-dependent | $v(t,i) = \frac{m\,i}{(K+i)}$ | $m = 7.62 \times 10^{14}$ [−, −]<br>$K = 1.74 \times 10^{15}$ [−, −] | $\text{hr}^{-1} \cdot \text{infected cell}^{-1}$<br>virions | 231 | 64.0 | $m$ = max production rate<br>$K$ = viral input at half max rate |
| **Time-dependent production rate models** | **Viral production rate** $\left(\mathbf{hr}^{-1} \cdot \textbf{infected cell}^{-1}\right)$ | **Parameter estimates (95% CI)** | **Units** | **RSS** | **$\Delta$AIC** | **Parameter legend** |
| Input-independent | $v(t,i) = \begin{cases} l\,(t-s) & , t > s \\ 0 & , \text{otherwise} \end{cases}$ | $l = 2.52$ [1.70, 3.75]<br>$s = 5.26$ [4.82, 5.75] | $\text{hr}^{-2} \cdot \text{infected cell}^{-1}$<br>hr | 98.1 | 26.3 | $l$ = slope of production rate<br>$s$ = time that production starts |
| Linear input-dependent | $v(t,i) = \begin{cases} r\,i\,(t-s) & , t > s \\ 0 & , \text{otherwise} \end{cases}$ | $r = 0.812$ [0.601, 1.10]<br>$s = 5.28$ [4.95, 5.62] | $\text{hr}^{-2} \cdot \text{virion}^{-1} \cdot \text{infected cell}^{-1}$<br>hr | 53.9 | 0 | $r$ = slope of per virion production rate<br>$s$ = time that production starts |
| Saturating input-dependent | $v(t,i) = \begin{cases} \frac{m\,i}{(K+i)}\,(t-s) & , t > s \\ 0 & , \text{otherwise} \end{cases}$ | $m = 2.68 \times 10^{15}$ [−, −]<br>$K = 3.30 \times 10^{15}$ [−, −]<br>$s = 5.27$ [5.00, 5.57] | $\text{hr}^{-2} \cdot \text{infected cell}^{-1}$<br>virions<br>hr | 53.9 | 2.00 | $m$ = max slope of production rate<br>$K$ = viral input at half max slope<br>$s$ = time that production starts |

**Table S3. Fits of viral production rate models to MDCK cell data.**

Rows correspond to distinct viral production rate models. Parameter estimates are given along with 95 percent confidence intervals for viral production rate model parameters. The model that is most supported by the data has $\Delta$AIC = 0, and models with higher $\Delta$AIC have less statistical support. Confidence intervals for high parameter estimates were omitted (see methods).

### Table S4. Fits of viral production rate models to A549 cell data.

| Time-independent production rate models | Viral production rate $\left(\mathrm{hr}^{-1} \cdot \text{infected cell}^{-1}\right)$ | Parameter estimates (95% CI) | | Units | RSS | ΔAIC | Parameter legend |
|---|---|---|---|---|---|---|---|
| Input-independent | $v(t,i) = l$ | $l = 0.158$ | $[0.086, 0.291]$ | $\mathrm{hr}^{-1} \cdot \text{infected cell}^{-1}$ | 187 | 61.5 | $l =$ viral production rate |
| Linear input-dependent | $v(t,i) = r\,i$ | $r = 0.0516$ | $[0.0276, 0.0965]$ | $\mathrm{hr}^{-1} \cdot \text{virion}^{-1} \cdot \text{infected cell}^{-1}$ | 198 | 63.9 | $r =$ per virion viral production rate |
| Saturating input-dependent | $v(t,i) = \frac{m\,i}{(K+i)}$ | $m = 0.326$<br>$K = 2.13$ | $[0.0266, 3.98]$<br>$[0.00878, 516]$ | $\mathrm{hr}^{-1} \cdot \text{infected cell}^{-1}$<br>virions | 185 | 63.0 | $m =$ max viral production rate<br>$K =$ viral input at half max rate |
| **Time-dependent production rate models** | **Viral production rate $\left(\mathrm{hr}^{-1} \cdot \text{infected cell}^{-1}\right)$** | **Parameter estimates (95% CI)** | | **Units** | **RSS** | **ΔAIC** | **Parameter legend** |
| Input-independent | $v(t,i) = \begin{cases} l\,(t-s) & , t > s \\ 0 & , \text{otherwise} \end{cases}$ | $l = 0.213$<br>$s = 5.00$ | $[0.133, 0.343]$<br>$[4.71, 5.31]$ | $\mathrm{hr}^{-2} \cdot \text{infected cell}^{-1}$<br>hr | 44.2 | 0 | $l =$ slope of viral production rate<br>$s =$ time that viral production starts |
| Linear input-dependent | $v(t,i) = \begin{cases} r\,i\,(t-s) & , t > s \\ 0 & , \text{otherwise} \end{cases}$ | $r = 0.0691$<br>$s = 4.99$ | $[0.0329, 0.0958]$<br>$[4.66, 5.33]$ | $\mathrm{hr}^{-2} \cdot \text{virion}^{-1} \cdot \text{infected cell}^{-1}$<br>hr | 55.7 | 10.2 | $r =$ slope of per virion viral production rate<br>$s =$ time that viral production starts |
| Saturating input-dependent | $v(t,i) = \begin{cases} \frac{m\,i}{(K+i)}\,(t-s) & , t > s \\ 0 & , \text{otherwise} \end{cases}$ | $m = 0.417$<br>$K = 1.91$<br>$s = 5.00$ | $[0.239, 0.730]$<br>$[0.703, 5.21]$<br>$[4.74, 5.28]$ | $\mathrm{hr}^{-2} \cdot \text{infected cell}^{-1}$<br>virions<br>hr | 42.4 | 0.232 | $m =$ max slope of viral production rate<br>$K =$ viral input at half max slope<br>$s =$ time that viral production starts |

**Table S4. Fits of viral production rate models to A549 cell data.**
Rows correspond to distinct viral production rate models. Parameter estimates are given along with 95 percent confidence intervals for viral production rate model parameters. The model that is most supported by the data has ΔAIC = 0, and models with higher ΔAIC have less statistical support.

**Table S5. Descriptions of interferon induction models.**

| Induction rate model | IFN induction rate $\left(\text{infected cell}^{-1}\right)$ | Parameter legend | Units |
|---|---|---|---|
| 1. Input-independent | $\lambda(i) = 1 + l$ | $l$ = rate of infected cell IFN induction | infected cell$^{-1}$ |
| 2. Linear input-dependent | $\lambda(i) = 1 + r\,i$ | $r$ = per virion rate of infected cell IFN induction | virion$^{-1}$ $\cdot$ infected cell$^{-1}$ |
| 3. Saturating input-dependent | $\lambda(i) = 1 + \frac{m\,i}{(K+i)}$ | $m$ = max rate of infected cell IFN induction<br>$K$ = viral input at half max rate | infected cell$^{-1}$<br>virions |

**Table S5. Descriptions of interferon induction models.** We considered three IFN induction models in which the induction rates are independent of time but differ based on the viral input. We fit an input-independent, linear input-dependent, and saturating input-dependent induction rate models to data at 8 and 18 hpi, giving a total of nine model combinations (see Table S6).

**Table S6. Fits of IFN induction models to IFNL1 data in A549 cells.**

| Model (0-8 hpi) | Parameter estimates (95% CI) | Model (8-18 hpi) | Parameter estimates (95% CI) | RSS | $\Delta$AIC |
|---|---|---|---|---|---|
| 1 | $l = 7.28$ $[3.20, 16.6]$ | 1 | $l = 790$ $[628, 994]$ | 110 | 19.2 |
| 1 | $l = 7.36$ $[3.24, 16.7]$ | 2 | $r = 211$ $[145, 309]$ | 128 | 25.5 |
| 1 | $l = 7.28$ $[3.20, 16.6]$ | 3 | $m = 1570$ $[994, 2480]$<br>$K = 2.10$ $[0.80, 5.48]$ | 107 | 20.2 |
| 2 | $r = 1.78$ $[0.87, 3.65]$ | 1 | $l = 782$ $[623, 983]$ | 68.8 | 0 |
| 2 | $r = 1.77$ $[0.86, 3.63]$ | 2 | $r = 215$ $[147, 314]$ | 88.1 | 10.1 |
| 2 | $r = 1.77$ $[0.86, 3.63]$ | 3 | $m = 1,510$ $[943, 2, 410]$<br>$K = 1.91$ $[0.70, 5.19]$ | 66.5 | 0.582 |
| 3 | $m = 2.08 \times 10^{16}$ $[-, -]$<br>$K = 1.17 \times 10^{16}$ $[-, -]$ | 1 | $l = 782$ $[623, 983]$ | 68.8 | 2.00 |
| 3 | $m = 1.11 \times 10^{16}$ $[-, -]$<br>$K = 6.28 \times 10^{15}$ $[-, -]$ | 2 | $r = 215$ $[147, 314]$ | 88.1 | 12.1 |
| 3 | $m = 6.18 \times 10^{15}$ $[-, -]$<br>$K = 3.49 \times 10^{15}$ $[-, -]$ | 3 | $m = 1,510$ $[943, 2, 410]$<br>$K = 1.91$ $[0.701, 5.19]$ | 66.5 | 2.58 |

**Table S6. Fits of IFN induction models to IFNL1 data in A549 cells.** Rows correspond to different combinations of the IFN induction rate models listed in Table 5 from 0-8 hpi and from 8-18 hpi. The first column gives the model number from 0-8 hpi, and the third column gives the model number from 8-18 hpi. Parameter estimates are given along with 95 percent confidence intervals for IFN induction rate model parameters. The model that is most supported by the data has ΔAIC = 0, and models with higher ΔAIC have less statistical support. Confidence intervals for high parameter estimates were omitted (see methods).

**Table S7. Fits of superinfection exclusion models to FACs data in MDCK cells.**

| Susceptibility model | Susceptibility of previously infected cells | Parameter estimates | RSS | $\Delta$AIC | Parameter legend |
|---|---|---|---|---|---|
| Input-independent | $0 < s < 1$ | $s = 0.0361$ $[0.0227, 0.0576]$ | 3,630 | 22.0 | $s$ = susceptibility of previously infected cells (1 being fully susceptible) |
| Input-dependent | $0 < r^i < 1$, for $i = 1, 2, 3, \ldots$ | $r = 0.293$ $[0.214, 0.402]$ | 2,230 | 0 | $r$ = per virion susceptibility of previously infected cells |

**Table S7. Fits of superinfection exclusion models to FACs data in MDCK cells.**
Rows correspond to distinct viral production rate models. The first model assumed that all rH3N1-infected cells had the same reduced probability of becoming infected with rH1N2 (input-independent). The second model assumed that the probability of being infected with rH1N2 decreased with cellular rH3N1 MOI (input-dependent). Parameter estimates are given along with 95 percent confidence intervals for viral production rate model parameters. The model that is most supported by the data has ΔAIC = 0, and models with higher ΔAIC have less statistical support.

**Table S8. RT-qPCR primers for the quantification of interferons, interferon stimulated genes, and endogenous control for MDCK (canine) and A549 (human) cell lines.**

| Canine Target | Chemistry | Forward Primer (5' to 3') | Reverse Primer (5' to 3') | Source |
|---|---|---|---|---|
| IFNB1 | SYBR Green | CTTGGATTCCTACAAAGAAGCAGC | TCCTCCTTCTGGAACTGCTGCA | OriGene (Rockville, MD, USA) |
| IFNL1 | SYBR Green | AACTGGGAAGGGCTGCCACATT | GGAAGACAGGAGAGCTGCAACT | OriGene (Rockville, MD, USA) |
| ISG15 | SYBR Green | GGTGGACAAATGCGACGAAC | ATGCTGGTGGAGGCCCTTAG | Pitha-Rowe, I et al. 2004 |
| Mx1 | SYBR Green | GGCTGTTTACCAGACTCCGACA | CACAAAGCCTGGCAGCTCTCTA | OriGene (Rockville, MD, USA) |
| β-actin | SYBR Green | GCGGGAAATCGTGCGTGACA | AAGGAAGGCTGGAAGAGTGC | Pitha-Rowe, I et al. 2004 |
| Human Target | Chemistry | Forward Primer (5' to 3') | Reverse Primer (5' to 3') | Source |
| IFNB1 | TaqMan FAM-MGB | N/A | N/A | Thermo Fisher Scientific (Waltham, MA, USA) |
| IFNL1 | TaqMan FAM-MGB | N/A | N/A | Thermo Fisher Scientific (Waltham, MA, USA) |
| GADPH | TaqMan JOE-TAMRA | N/A | N/A | Thermo Fisher Scientific (Waltham, MA, USA) |
| ISG15 | SYBR Green | TCTGTGCCCCTGGAGGACTTGA | TGCTGCTTCAGCTCTGATGCCA | Fan, W et al. 2014 |
| Mx1 | SYBR Green | GAATCCTGTACCCAATCATGTG | TACCTTCTCCTCATATTGGCT | Fan, W et al. 2014 |
| β-actin | SYBR Green | CGAGACATTCAACACCCCAAC | AGCCAGGTCCAGACGCAAG | Fan, W et al. 2014 |

**Table S8. RT-qPCR primers for the quantification of interferons, interferon stimulated genes, and endogenous control for MDCK (canine) and A549 (human) cell lines.** SYBR green primers were used for the quantification of canine targets of IFNB1, IFNL1, ISG15, and Mx1, with ß-actin as the endogenous control. For A549, Taqman assays were used for the quantification of IFNB1 and IFNL1 with GADPH as the endogenous control and SYBR green chemistry was used for the quantification of ISG15 and Mx1, with ß-actin as the endogenous control.
